# HyGAnno: Hybrid graph neural network-based cell type annotation for single-cell ATAC sequencing data

**DOI:** 10.1101/2023.11.29.569114

**Authors:** Weihang Zhang, Yang Cui, Martin Loza, Sung-Joon Park, Kenta Nakai

**Affiliations:** Department of Computational Biology and Medical Sciences, Graduate school of Frontier Sciences, University of Tokyo, 5-1-5 Kashiwanoha, Kashiwa, 277-8561, Japan; Human Genome Center, Institute of Medical Science, University of Tokyo, 4-6-1 Shirokaneda, Tokyo, 108-8639,Japan

**Keywords:** single-cell ATAC sequencing, label transfer learning, single-cell multi-omics, graph embedding

## Abstract

Reliable cell type annotations are crucial for investigating cellular heterogeneity in single-cell omics data. Although various computational approaches have been proposed for single-cell RNA sequencing (scRNA-seq) annotation, high-quality cell labels are still lacking in single-cell ATAC sequencing (scATAC-seq) data, because of extreme sparsity and inconsistent chromatin accessibility between datasets. This calls for novel cell type annotation methods in scATAC-seq, to better explore cell type-specific gene regulatory mechanisms and provide a complementary epigenomic layer to scRNA-seq data. Here, we present a novel automated cell annotation method that transfers cell type information from a well-labeled scRNA-seq reference to an unlabeled scATAC-seq target, via a parallel graph neural network, in a semi-supervised manner. Unlike existing methods that utilize only gene expression or gene activity features, HyGAnno integrates genomewide accessibility peak features to facilitate the training process. In addition, HyGAnno reconstructs a reference-target cell graph that can be used to detect cells with low prediction reliability, according to their specific graph connectivity patterns. HyGAnno was tested using large datasets and demonstrated the advantages of accurate cell annotation, interpretable cell embedding, robustness to noisy reference data, and adaptability to tumor tissues.

## 1 Background

Single-cell sequencing technologies have become increasingly popular in contemporary biology and promote the understanding of the molecular mechanisms underlying the formation of heterogeneous cell populations. In particular, multi-omics approaches coupled with chromatin accessibility using single-cell ATAC sequencing (scATAC-seq) and transcriptome analysis using single-cell RNA sequencing (scRNA-seq) are widely used to dissect gene regulation at the single-cell level. Recently, many studies have achieved computational and experimental improvements, thereby providing new possibilities for single-cell multi-omics analyses [1, 2].

A fundamental issue in using single-cell multi-omics datasets is that populations detected by means of omics sequencing must be annotated and invariably matched, which is a prerequisite for precisely integrating multi-omics features. While cell annotation has been considered in the context of scRNA-seq [3–5], it still presents challenges in terms of other single-cell datasets. Specifically, given the increasing availability of public scATAC-seq datasets, there is a high need for cell annotations based on chromatin accessibility [6, 7].

To overcome the paucity of scATAC-seq cell annotation, many studies have employed the following three strategies, with room for improvement: (1) Mapping peak enrichment to important genes. However, owing to the high sparsity of ATAC peaks and incomplete establishment of libraries, the annotations obtained thus are of very low resolution and easily overlap, leading to imprecise annotations; (2) Incorporating the cell labels of scRNA-seq data as references, such as Seurat [8], scJoint [9], scGCN [10], and Conos [11]. For convenience, the genome-wide accessible peak (peak-level) features of the target scATAC-seq are transformed into gene activity (gene-level) features [12, 13]. Then cell labels in reference data are transferred to the target data according to their gene-level similarity. Although these tools have been widely used because of their simplicity, the inherent peak-level information is insufficiently utilized; (3) Incorporating the cell labels of scATAC-seq data as references, such as Cellcano [14] and EpiAnno [15]. Cellcano converts both the reference and target scATAC-seq data into gene activity matrices, resulting the information loss problem of the peak-level features. Although EpiAnno uses peak-level features within a Bayesian network, the availability of well-annotated scATAC-seq datasets that can serve as references is limited, potentially constraining its efficiency.

This study provides a novel graph-based deep learning approach called HyGAnno to predict cell types in scATAC-seq experiments. Our method represents the cell populations in scRNA-seq and scATAC-seq datasets as undirected graphs and links the graphs using anchor cells. These graphs are then embedded and learned using graph neural networks, to transfer the cell labels of the scRNA-seq data to the corresponding cell populations in the scATAC-seq data. This approach not only ensures precise annotations by training models with genome-wide accessibility peak information, but also reconstructs target-reference cell graphs, which can describe the prediction reliability.

## 2 Results

### 2.1 The HyGAnno framework

HyGAnno predicts cell labels for scATAC-seq data by aligning the embeddings of the same ATAC cells viewed in different graphs (Fig. 1). First, HyGAnno builds a hybrid graph by computing the similarity of gene expression and gene activity features (collectively termed gene-level features) between RNA cells and ATAC cells. ATAC cells showing similar gene-level similarity with RNA cells (termed ATAC anchor cells) remain in the hybrid graph, whereas non-ATAC anchor cells are removed from the graph. Similarly, an ATAC graph can be constructed by computing the similarity of genome-wide chromatin accessible peak features (termed peak-level features) among ATAC cells. Then, HyGAnno employs parallel graph neural networks to embed hybrid and ATAC graphs into separate latent spaces and minimizes the distance between the embeddings of the same ATAC anchor cells in the two graphs. Using this approach, cell labels can be automatically transferred from scRNA-seq data to scATAC-seq data. In addition, HyGAnno reconstructs a consolidated reference-target cell graph that shows more complex graph structures, thus inspiring us to describe ambiguous predictions based on abnormal target-reference cell connections. Besides ensuring precise cell annotations, HyGAnno is also expected to outperform existing methods in visualizing interpretable cell clusters, handling noise cell labels in reference data, and discovering cell type-specific peaks.

**Figure 1.**
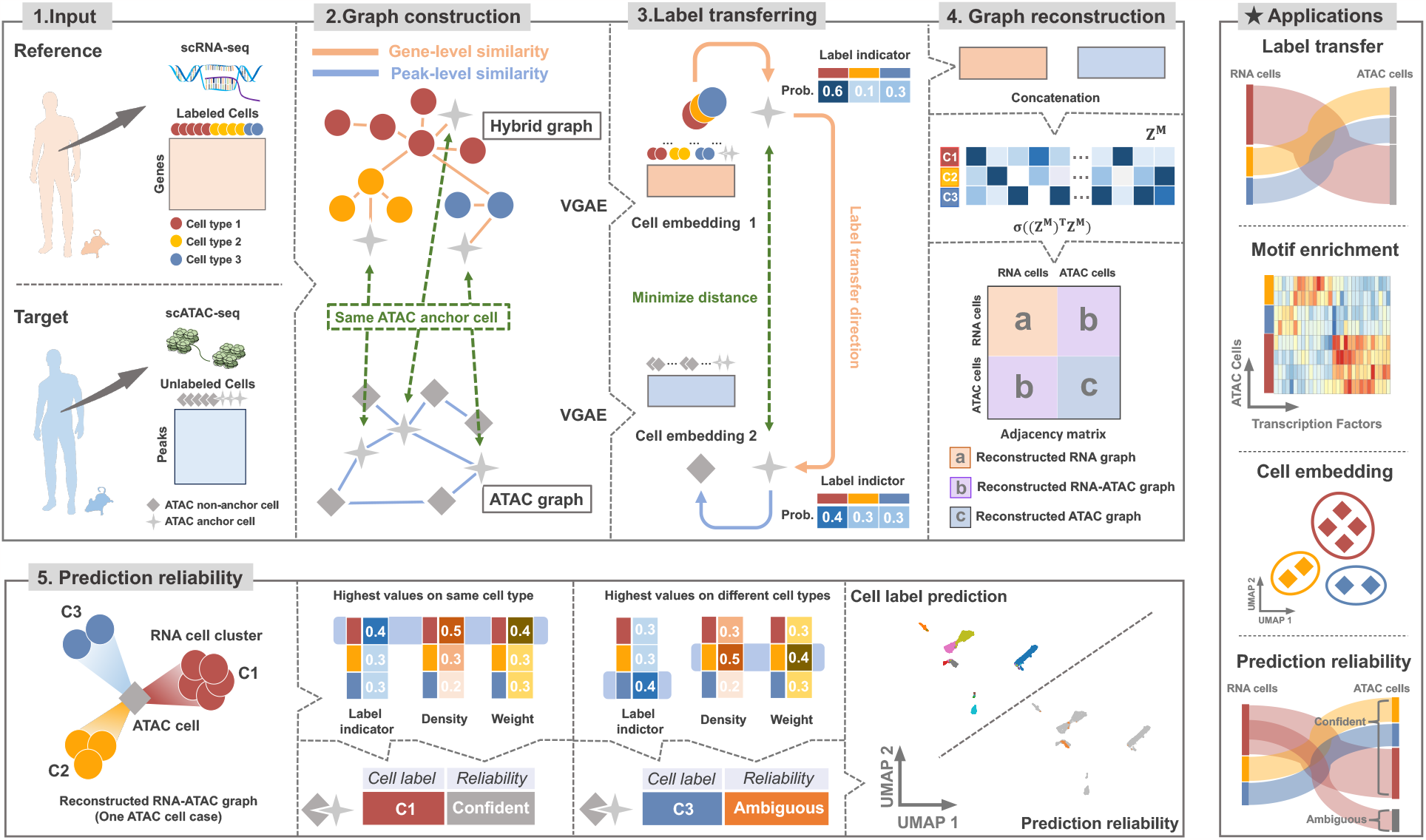
Overview of the HyGAnno framework. HyGAnno adopts graph neural networks to transfer cell label information from labeled scRNA-seq reference data to unlabeled scATAC-seq target data. The input of HyGAnno consists of an unlabeled peak matrix from scATAC-seq data and a well-labeled gene expression matrix from scRNA-seq data. HyGAnno is designed to train a model with both gene-level and peak-level features. For gene-level features, HyGAnno constructs a hybrid graph containing labeled cells in scRNA-seq (termed RNA cells) and cells highly correlated with RNA cells in scATAC-seq (termed ATAC anchor cells). For peak-level features, an ATAC graph is constructed containing all cells in scATAC-seq (termed ATAC cells). Then, HyGAnno embeds graphs with parallel variational graph auto-encoders and aligns the embedding of anchor ATAC cells viewed by two graphs. In this process, the cell labels are transferred from the RNA cells to ATAC cells. Meanwhile, a more informative RNA-ATAC cell graph can be constructed. When an ATAC cell in the reconstructed graph shows the highest connectivity to the cell type consistent with the label indictor, the prediction reliability is confident, otherwise, ambiguous. Prob., probability; TF, transcription factor; VGAE, variational graph auto-encoder.

### 2.2 Comparative annotation performance of HyGAnno with benchmarking methods

We first benchmarked HyGAnno against Seurat [8], scJoint [9], scGCN [10], and Conos [11], which utilized the cell labels of the scRNA-seq data. These methods were applied to mouse brain and lung datasets, as well as human peripheral blood mononuclear cell (PBMC) and bone marrow mononuclear cell (BMMC) datasets. Performance was measured in terms of Accuracy (ACC) and Normalized Mutual Information (NMI). Overall, HyGAnno achieved the highest average ACC (0.87) among the four datasets (Fig. 2a), followed by Seurat (0.77) and scJoint (0.73). Even though scGCN also takes advantage of graph convolutional network (GCN), the ACC and NMI were significantly lower than HyGAnno because it trains model without reference graph.

**Figure 2.**
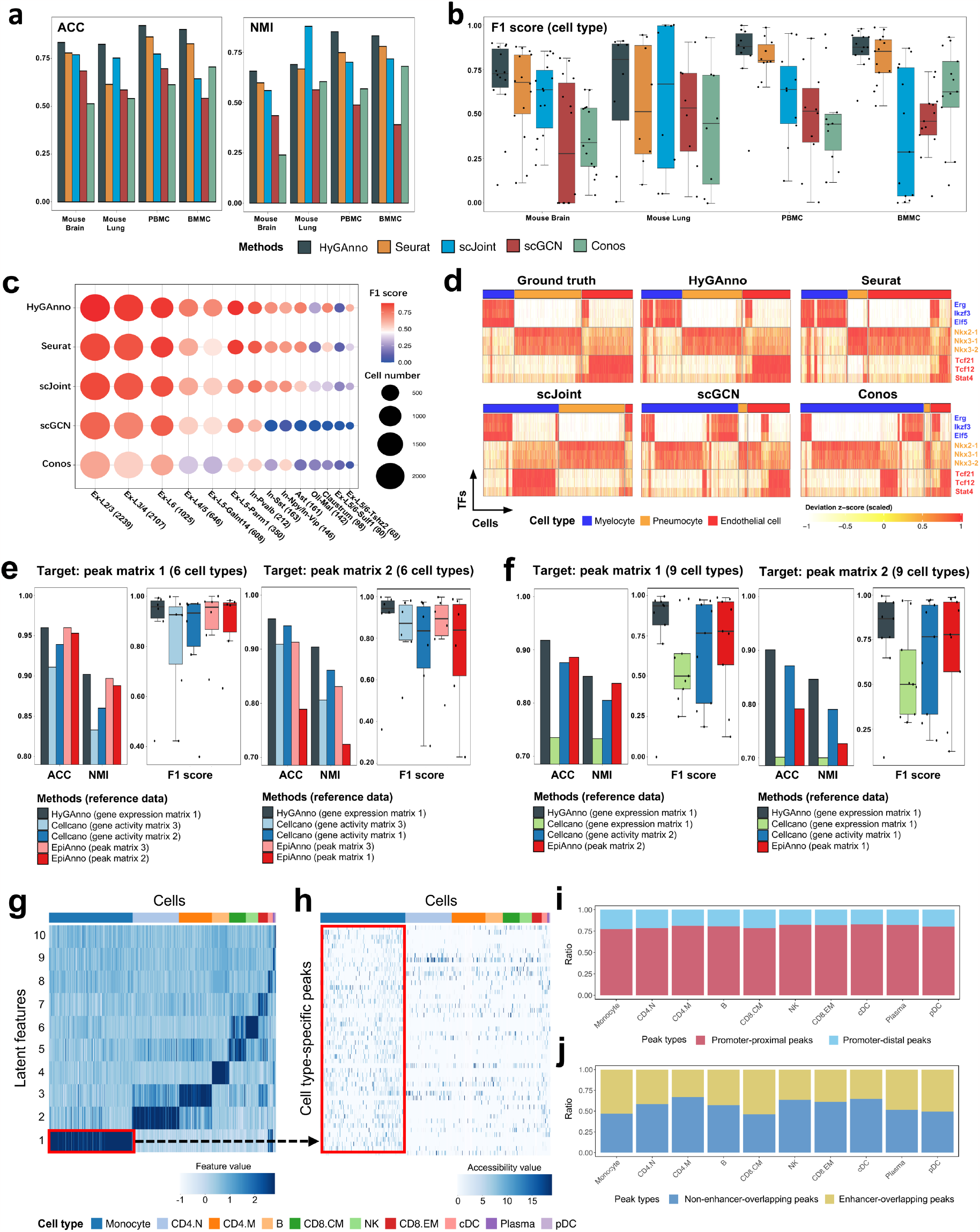
Comparative analysis of HyGAnno with RNA-referenced and ATAC-referenced benchmarks. **(a)** General cell annotation comparisons of HyGAnno, Seurat, scJoint, scGCN, and Conos, evaluated in terms of ACC and NMI, on four target scATAC-seq datasets of mouse brain (8,055 cells with 14 types), mouse lung (7,499 cells with 8 types), PBMC (7,828 cells with 10 cell types) and BMMC (14,753 cells with 13 types). **(b)** Annotation performance for each cell type evaluated in terms of F1 scores. Inside the boxes, dot points represent the F1 scores for different cell types. The top, middle, and bottom lines of the box mark the 75th, 50th, and 25th percentiles, respectively. Whiskers extend to data points within 1.5 times the interquartile range. **(c)** Comparisons of F1 scores regarding cell number of each cell type in mouse brain data. The brackets indicate the cell number of each cell type. **(d)** TF motif activity scores calculated using chromVAR, based on the prediction result of each benchmark. **(e)** Cell type prediction comparisons of HyGAnno, Cellcano and EpiAnno evaluated in terms of the ACC, NMI and F1 score. Two peak matrices containing 2,588 cells and 3,570 cells were used as the target. Six broad cell types were used for evaluation. **(f)** Same target data with nine cell types. **(g, h)** Heatmap of the latent space and the peak count matrix, where rows are cell type-specific features and cell type-specific peaks, respectively; columns are cells colored by ground truth. **(i)** Cell type-specific peak annotation based on their genome locations. **(j)** Overlapped cell type-specific enhancers with prompter-distal peaks. ACC, Accuracy; NMI, Normalized Mutual Information; PBMC, peripheral bone mononuclear cell; BMMC, bone marrow mononuclear cell

To better assess the performance of the imbalanced datasets, we calculated the F1 scores for each cell type according to different benchmarks (Fig. 2b). HyGAnno achieved a better performance for most cell types across the four datasets. In particular, we confirmed that HyGAnno had a higher predictive resolution for rare cell types (Fig. 2c). For example, Claustrum and Ex-L5/6-Tshz2 in mouse brain data contained only tens of cells. In addition, even when the cell number of the reference (2623) was twice as small as that of the target (7499) in the mouse lung data, unlike other methods that struggle with insufficient training, HyGAnno precisely annotated the cell types, e.g., endothelial cells (Supplementary Fig. 1). Then, to investigate how cell labelling influences downstream analyses, we analyzed the enrichment of transcription factors (TFs) in the accessibility peaks of the predicted ATAC cell types in mouse lung, using the deviation of z-scores calculated by chromVAR [16]. We assessed the enrichment patterns of the cell type-specific TFs, such as, *Erg, Nkx2-1*, and *Tcf21* [17–20] based on the ground truth and predicted cell labels of myelocytes, pneumocytes, and endothelial cells. The TF motif enrichment patterns based on the predictions of HyGAnno were highly similar to those of the ground truth (Fig. 2d), which highlights the importance of appropriate annotation to ensure accurate downstream analysis.

Next, for a comprehensive comparison, we benchmarked HyGAnno with Cellcano [14] and EpiAnno [15], which utilize the cell labels of the scATAC-seq data. To assess the prediction accuracy of HyGAnno, Cellcano, and EpiAnno, we used fixed target data, but different reference data. First, we checked the annotation results for six cell types in scATAC-seq PBMC data. HyGAnno performed similar performance to EpiAnno and Cellcano (Fig. 2e). Then, to assess the annotation ability of more cell types, we increased the number of cell types to nine, including more subtypes from the original research [21]. We observed pronounced superiority of HyGAnno in achieving high-resolution cell annotation, with a median of F1 scores of 0.86, much higher than those for both Cellcano and EpiAnno (0.75) (Fig. 2f). Finally, to examine whether the performance of Cellcano could be improved, we trained Cellcano using the same scRNA-seq input as HyGAnno. However, the results worsened (indicated by green boxes in Fig. 2f), suggesting that HyGAnno was more capable of exploiting high-quality reference data.

As HyGAnno takes genome-wide accessible peak (peak-level) features as the input (Supplementary Fig. 2), we can extract the optimized weight matrices from deep neural networks to interpret the importance of each peak-level features on cell type identification. We trained a one-layer network for PBMC data, where cell type-specific latent features (Fig. 2g) are directly connected with the input peak-level features. This configuration enables the efficient identification of cell type-specific peaks (Fig. 2h). Given that the cellular specificity is largely governed by cis-regulatory elements (CREs) across entire genome sequences, we annotated cell type-specific peaks according to their proximity to transcription start sites (Fig. 2i). We found that around 25% of these peaks were promoter-distal, indicating the importance of distal intergenic regions in distinguishing cell types. To further investigate the CREs in these promoter-distal peaks, we mapped them to scEnhancer [22], a single-cell enhancer database, and found that half of these peaks were overlapped with cell type-specific enhancers (Fig. 2j). This result proved the powerful ability of HyGAnno in extracting cell type-specific characteristics by giving weights to not only the promoter-proximal regions but also the CREs in distal intergenic regions, thus guaranteeing the precision of the high-resolution cell-type annotation.

### 2.3 Leveraging peak-level features improves cell embeddings

To investigate the effect of peak-level information on cell embeddings, we generated cell embeddings using five methods: HyGAnno, Latent Semantic Indexing (LSI) [13], Principle Component Analysis (PCA), scJoint, and scGCN. We assessed these methods by calculating the Silhouette Width (SW) scores for the embedding spaces across various datasets (Fig. 3a). Notably, HyGAnno outperformed PCA, scJoint, and scGCN, all of which employ only gene-level features. The small interquartile range of HyGAnno indicates its high robustness. LSI also considers peak-level features; however, its performance is limited owing to the lack of scRNA-seq reference data. To interpret the cell embeddings, we then visualized the cell embeddings of the PBMC data across different methods using Uniform Manifold Approximation and Projection (UMAP) [23] (Fig. 3b). Taking the T-cell subtypes as an example, we observed that HyGAnno, LSI, PCA, and scJoint could distinguish the four T cell subtypes (red circles in Fig. 3b) from B cells and monocytes. In contrast, scGCN embedded all T cell subtypes together, creating a mixture that resulted in the lowest average silhouette width (ASW) score. Both HyGAnno and LSI captured the differentiation processes from naïve CD4 T cells (CD4.N) to memory CD 4 T cells (CD4.M) [24]. However, HyGAnno achieved a higher ASW score (0.43) than LSI (Fig. 3b) and separated NK cell cluster by taking the cell embedding information from reference data (Supplementary Fig. 3a). Although scJoint detected the discrete cell islands, it exhibited a lower ASW score (0.24) than that of HyGAnno. This result indicated a misunderstanding of cell development and differentiation information led by the scJoint forcing the cells with the same predicted labels being near the random centers. To further demonstrate the strength of the cell embedding obtained from HyGAnno in cellular differentiation analysis, we calculated the pseudotime of CD4.N and CD4.M based on different cell embeddings (Fig. 3c) and found that only the cell embedding from HyGAnno can achieve continuous trajectory to show the differentiation process. In addition to the PBMC data, the UMAP visualization of BMMC also suggests the interpretable cell embeddings (Supplementary Fig. 3b, c); for instance, the differentiation trend from common lymphoid progenitors (CLPs) to Precursors of lymphocytes (Pre.B cells) and B cells [25] (Supplementary Fig. 3d). The highest ASW score (0.33) again indicates the capacity of HyGAnno to form cell clusters. Collectively, these results highlighted that the collaborative training of HyGAnno using both gene expression information from the scRNA-seq reference and genome-wide accessibility information from the scATAC-seq target, which provides interpretable cell embeddings, ensuring the trajectory analysis and describing cellular similarity and potential differentiation process.

**Figure 3.**
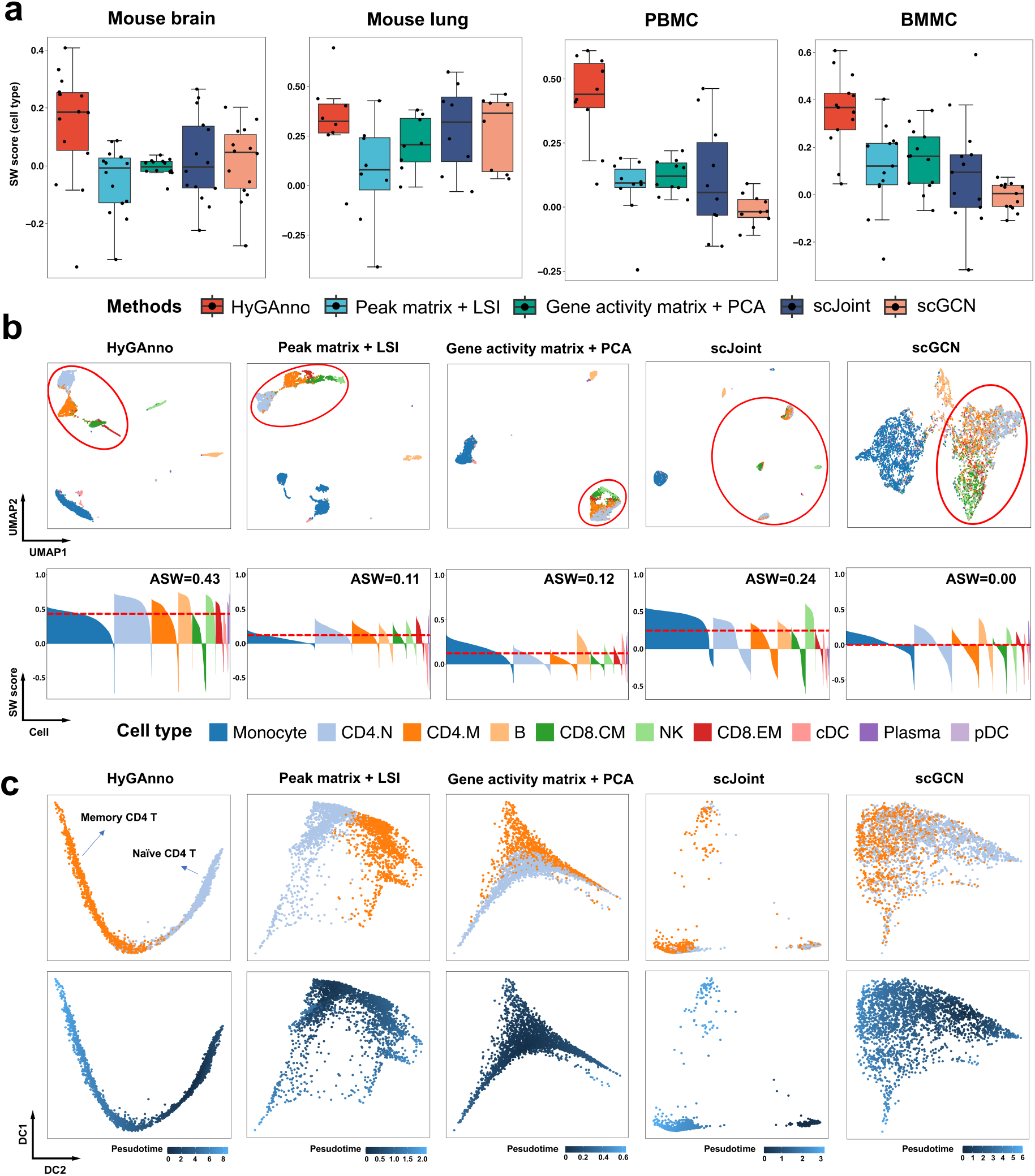
Evaluation of cell embeddings. **(a)** SW scores of the cell embeddings obtained using HyGAnno, LSI, PCA, scJoint, and scGCN. Inside the boxes, dot points represent SW scores for different cell types. The top, middle, and bottom lines of the box mark the 75th, 50th, and 25th percentiles, respectively. Whiskers extend to the data points that are within 1.5-times the interquartile range. **(b)** UMAP plots (upper row) and the corresponding SW score (bottom row) of PBMC cell embeddings. The cells are colored based on ground truth cell types. The four T cell subtypes are surrounded by red circles. **(c)** Trajectory analysis based on the cell embeddings of CD4.N cells and CD4.M cells. The cells are colored based on ground truth cell types (upper row) and pseudotime calculated using DPT (bottom row). UMAP, Uniform Manifold Approximation and Projection; SW, Silhouette Width; DPT, Diffusion Pseudotime

### 2.4 HyGAnno resists noisy cell labels in reference data

Most scRNA-seq data are manually annotated and adjusted by exploring the expression patterns of the marker genes, which inevitably leads to incorrect cell labels. To assess how noisy cell labels in scRNA-seq references affect the cell type annotation, we randomly generated ten false cell lists for each data, by increasing the noise level proportion (*p*) from 0.1 to 0.9. In each proportion, the average performance of the five methods across the four datasets were reported in terms of weighted F1 score, ACC, and NMI (Fig. 4a).

**Figure 4.**
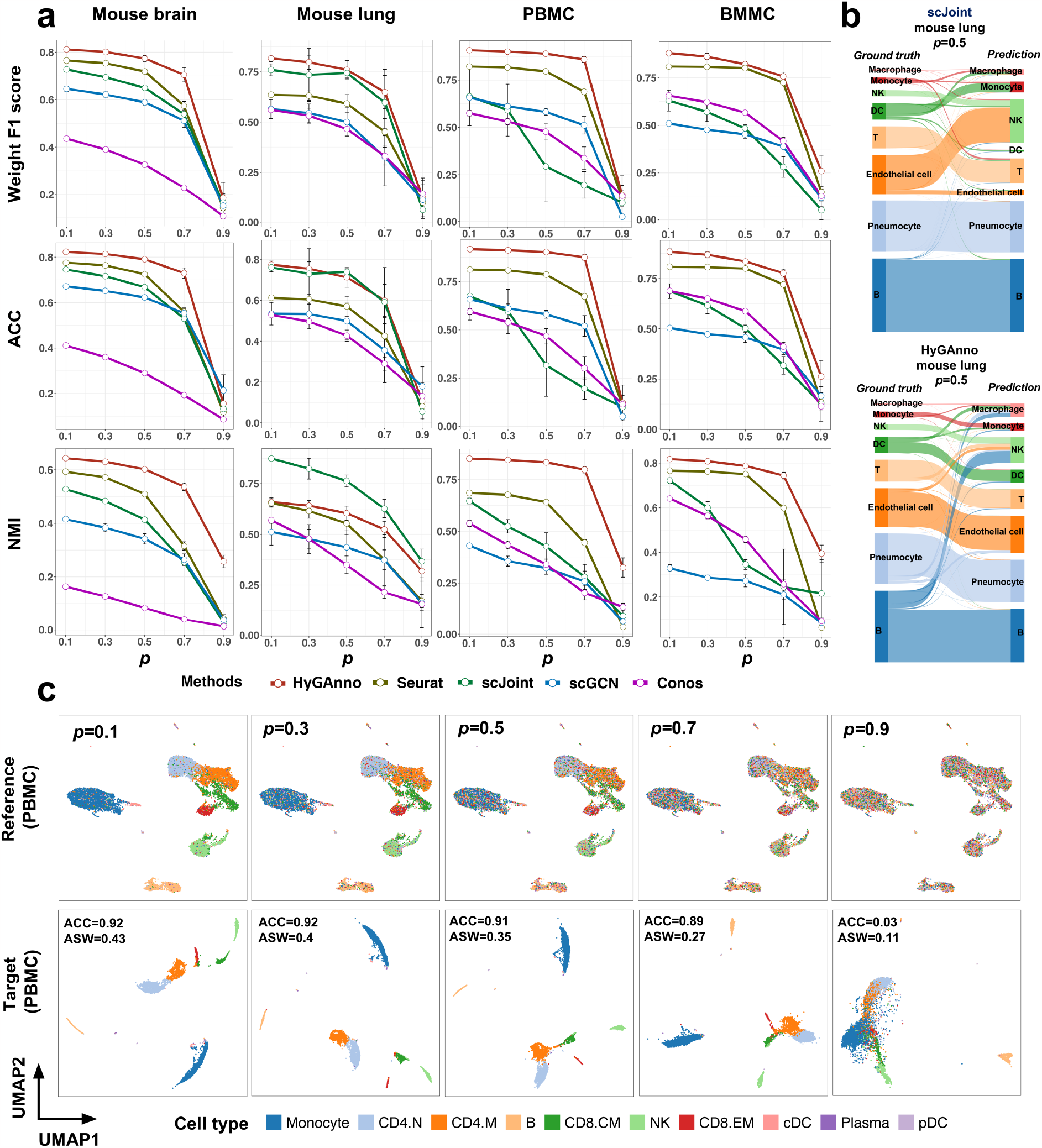
Evaluation of HyGAnno against noisy reference data. **(a)** Weighted F1 scores, ACC, and NMI were calculated for HyGAnno, Seurat, scJoint, scGCN, and Conos across four datasets with increasing noise level *p* from 0.1 to 0.9. Each point indicates the mean value among 10 randomly generated cell label lists and the whisker is extended by standard deviation. **(b)** Sankey plots indicating the details of how cell labels are transferred from reference to target data of HyGAnno and scJoint under the condition of *p* = 0.5 in mouse lung data. **(c)** UMAP plots of PBMC reference data colored by noise labels (upper row) and PBMC target data colored by predicted labels (bottom row), with *p* from 0.1 to 0.9.

We found that all methods deteriorated as *p* increased, and finally lost their annotation ability when *p* reached 0.9 (Fig. 4a). HyGAnno outperformed the other methods by consistently achieving the highest weighted F1 and ACC scores for all data. Meanwhile, Seurat was the second-best performer, with similar performance to HyGAnno in the mouse brain and BMMC data. In the mouse lung data, scJoint had higher NMI scores than HyGAnno. This discrepancy arises because a high NMI score is obtained if two samples of the same class are classified into the same group. Hence, a higher NMI paired with a relatively lower weighted F1 score indicates that scJoint might incorrectly annotate the entirety of some cell clusters. To verify this, we illustrated the prediction results of HyGAnno and scJoint by means of Sankey plots under the condition of *p* = 0.5 in mouse lung data (Fig. 4b). We observed that scJoint failed to annotate whole clusters of monocytes, dendritic cells, and endothelial cells, in which most endothelial cells were labeled as NK cells, despite the huge gap in biological functions between them. In contrast, although some B cells were wrongly labeled as NK cells by HyGAnno, the prediction was still acceptable, as there were no entirely mislabeled cases in any clusters. We also visualized the reference and target data of PBMC with p from 0.1 to 0.9 using UMAP (Fig. 4c). We found that before *p* reached 0.9, not only was the cell annotation accuracy of HyGAnno almost unchanged from 0.92 to 0.89, but the cell embedding structures was also preserved, with an ASW score of 0.27. All the above results indicate the power of HyGAnno to resist noisy cell labels in the reference data, thus lowering the dependence on perfectly annotated scRNA-seq data.

### 2.5 HyGAnno describes the reliability of predicted cell labels

To assess how HyGAnno handles ambiguous predictions, we first reconstructed reference-target cell graphs using HyGAnno from four datasets: mouse brain, mouse lung, PBMC, and BMMC. We quantified the differences between the reconstructed and initial graphs based on the density of edges (DLE) between RNA and ATAC cells. Based on the heatmaps shown in Supplementary Fig. 4, compared with that in the initial graph, reconstructed graph showed more non-diagonal DLE values, indicating that the reconstructed graphs could amplify the abnormal cell-cell connections hidden in the initial ones. This characteristic in the reconstructed graph inspired us to design the strategy mentioned in section 5.6 to detect ambiguous predictions. We found that the ambiguous predictions of the above four datasets are typically scattered within the regions with highly similar cell types (Fig. 5a); for instance, a clear cell boundary between the clusters of granulocyte-monocyte progenitors (GMP) and monocytes in the BMMC data. Because these ambiguously predicted cells may be unreliable, we eliminated them from the datasets and observed notable improvements across several evaluation metrics (Supplementary Fig. 5). This observation suggested the removal of the cells in target data with ambiguous predictions.

**Figure 5.**
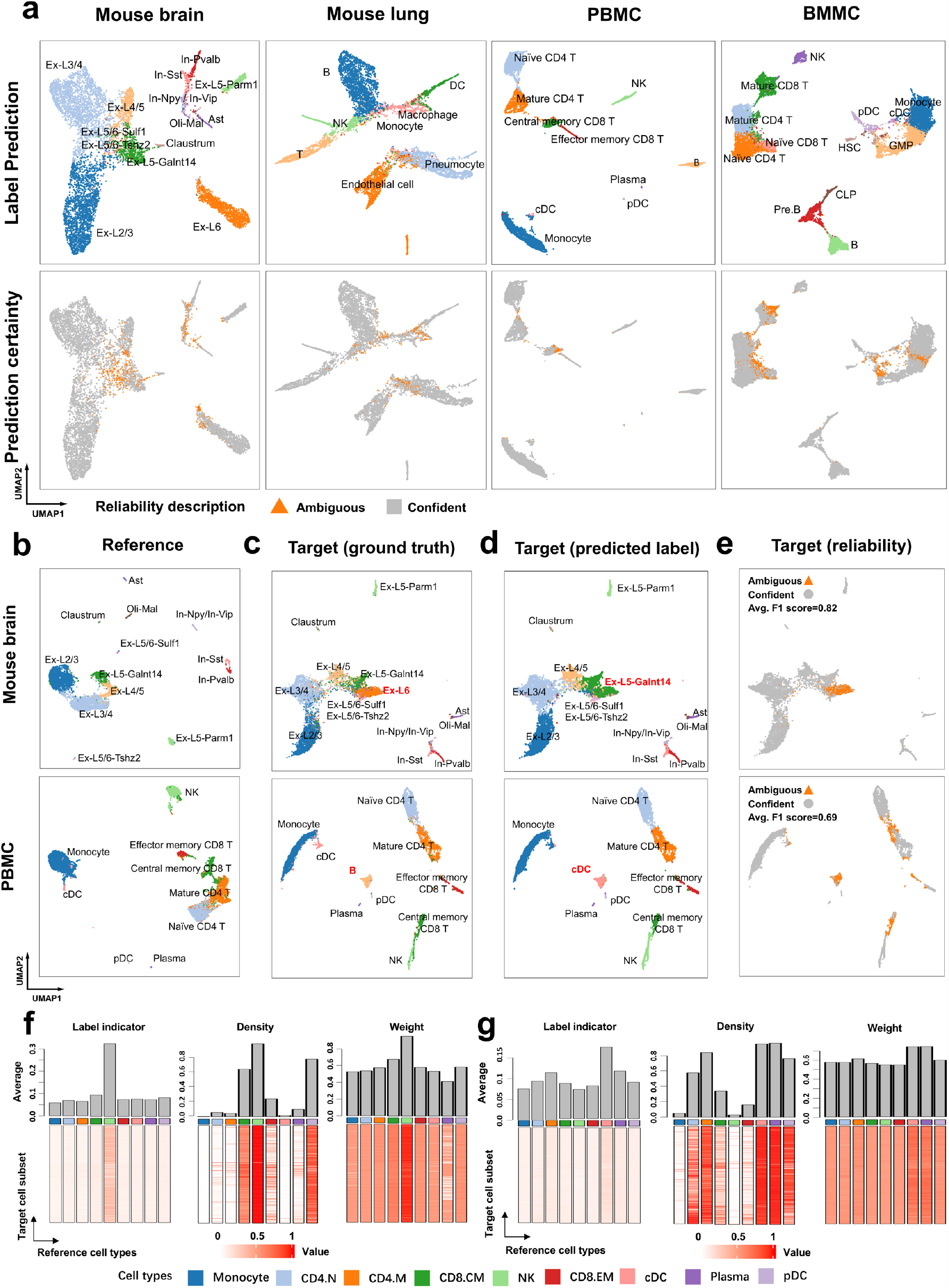
Description of prediction reliability based on the reconstructed graph. **(a)** Cell type annotation and the prediction reliability of four data. **(b)** UMAP plot of the reference data of mouse brain and PBMC, excluding the Ex-L6 cells and B cells, respectively. Cells are colored based on their ground truth labels. **(c, d, e)** UMAP plot of the target data of mouse brain and PBMC, including Ex-L6 cells and B cells. Cells are colored based on their ground truth labels, prediction labels, and prediction reliability, respectively. **(f, g)** Heatmaps of label indicator, density, and weight matrix obtained from the reconstructed graph of PBMC data.

However, the ambiguous predictions are caused not only by the high cell type similarity in the target data, but also by the cell-type bias between the reference data and target data. In addition to the detection of ambiguous predictions for the non-biased datasets mentioned above, we next simulated the scenarios of cell type-biased datasets. We manually removed Ex-L6-Tle4 cells and B cells from the reference mouse brain data and PBMC data, respectively (Fig. 5b), while the target data remained unchanged (Fig. 5c). After predicting the cell labels, we compared the results with the ground truth and found that most Ex-L6 cells and B cells were inappropriately labeled as Ex-L5-Galnt14 cells and cDC cells, respectively (Fig. 5d). However, the clusters of these two cell types were successfully captured by the ambiguous predictions, with high F1 scores of 0.82 and 0.69 (Fig. 5e), indicating the ability of HyGAnno to handle the cell type-insufficiency problem of reference data.

Finally, to further demonstrate how the prediction reliability is denoted, we used the above cell type-biased PBMC data as an example and visualized the label indicator, density, and weight matrices (Fig. 5f, g), which helped us formulate the pattern of the abnormal cell-cell connections amplified by the reconstructed graph. According to the label indicator matrix (Fig. 5f), the target cell subsets (rows) had the highest values for the NK cell type (column); hence, they were predicted to be NK cells. Based on the density and weight matrices, which summarize the connection patterns in the reconstructed graph (Supplementary Fig. 6a), we found that the target NK-like cells were densely connected to the reference NK cells. This consistent observation from two scopes suggests confident labeling prediction. In contrary, for another target cell subset (Fig. 5g), although most of them were predicted to be cDC, the inconsistent dense connection with not only the reference cDC, but also plasma, pDC, and CD4.M cells lowered their prediction reliability (Supplementary Fig. 6b). Collectively, as an additional procedure after HyGAnno cell annotation, the cell type prediction reliability can be described using the reconstructed graph. For user convenience, we packaged this function as a script and highly recommended the removal of cells with ambiguous descriptions.

### 2.6 HyGAnno annotates neoplastic and normal cells in breast cancer

Because of the shared lineages of tumor cells with their normal counterparts [26], it is challenging to identify them using marker genes or clustering analysis. To assess the capacity of HyGAnno to annotate cells within the tumor microenvironment (TME), we gathered scRNA-seq and scATAC-seq (Fig. 6a) as reference and target datasets from breast cancer and annotated them using the same approach outlined in the original works [27, 28], in which manual cell annotation failed to differentiate tumor and epithelial cells in the scATAC-seq data. To address this, we leveraged cell labels of scRNA-seq data using HyGAnno and obtained new cell annotations, wherein immune, fibroblast, and endothelial cells remained consistent with the manual annotation results, while partial epithelial cells were reannotated as tumor cells with confident prediction reliability (Fig. 6b). Because the ground truth of the scATAC-seq data was unclear, we presented the following evidence to support the reliability of the identified normal and tumor cells.

**Figure 6.**
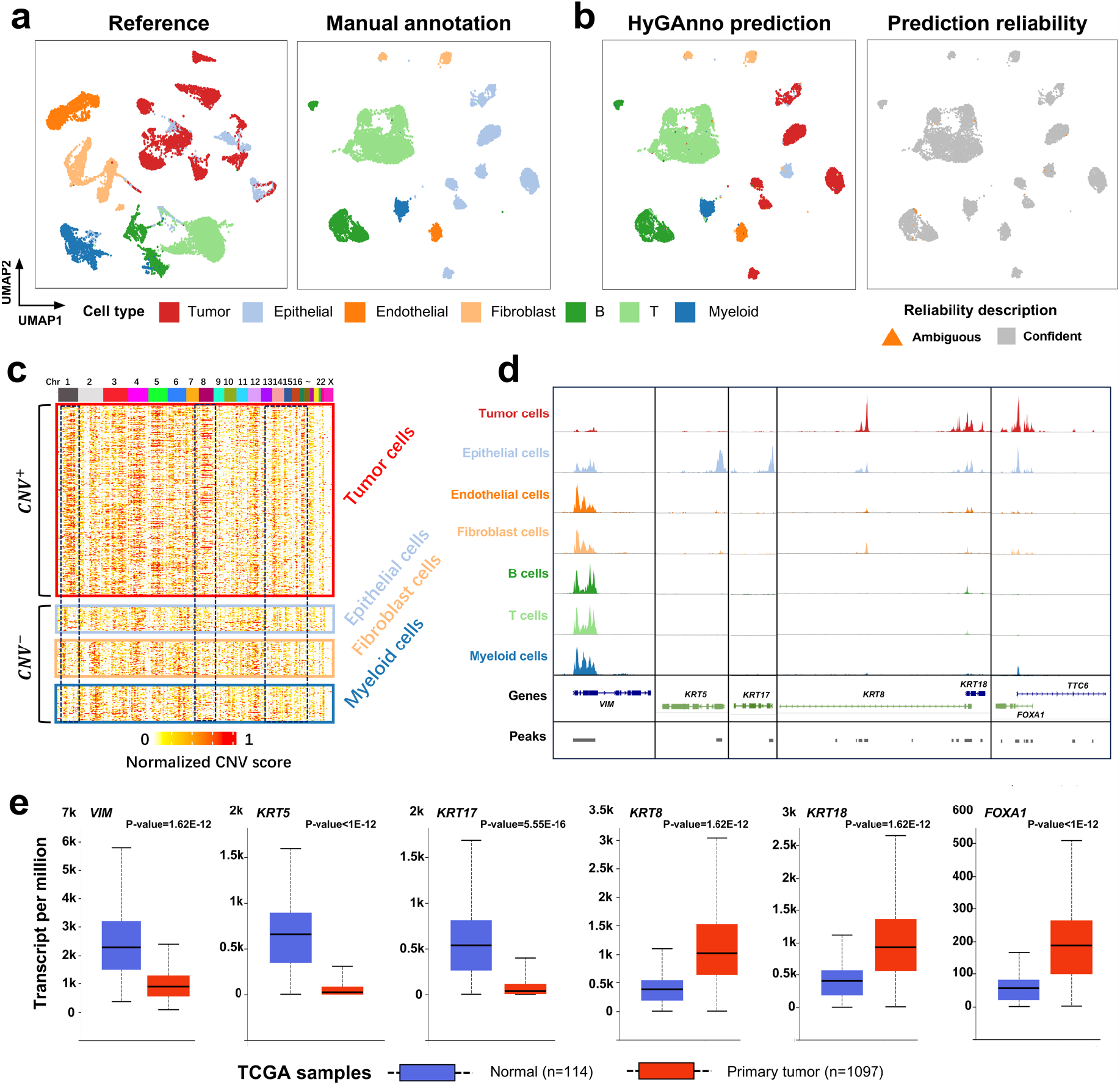
Evaluation of cell type prediction in the tumor microenvironment using HyGAnno. **(a)** UMAP plots of reference data and target data from breast cancer, colored based on manually annotated cell types. **(b)** UMAP plots of target data, colored based on HyGAnno cell type prediction and prediction reliability. **(c)** Normalized copy number variation patterns of each cell cluster. **(d)** Enrichment of differential accessibility peaks of tumor and normal cells in the promoter regions of potential signature genes. **(e)** Gene expression differences of observed genes between 1097 breast cancer and 114 normal samples.

Firstly, we demonstrated the differences in DNA amplification between predicted tumor and normal cells. Based on previous works [27, 29], we inferred the DNA copy number variation (CNV) signals by calculating the normalized CNV scores of the scATAC-seq data (Fig. 6c). We found that tumor cells have stronger CNV signals than the epithelial cells on chromosome 1, which has been reported to be involved in quantitative anomalies in 50% ∼ 60% of breast cancers [30]. Because malignant transformation occurs predominantly in breast tissues from epithelial cells [31], we used the CNV patterns of other cell types, such as the fibroblasts and the infiltrating myeloid cells, as the normal background. The predicted epithelial cells showed an amplification pattern similar to that of the normal background on each chromosome, indicating a high possibility of these cells being normal.

To further demonstrate the accuracy of the HyGAnno prediction, we extracted the differential accessibility peaks between tumor cells and epithelial cells, and checked whether they were enriched in the promoter regions of certain genes, as this might reflect the specific expression patterns of these two cell categories. The peaks of epithelial cells and tumor cells were notably enriched in the gene sets of *‘VIM, KRT5, KRT17’* and *‘KRT8, KRT18, FOXA1’*, respectively (Fig. 6d, Supplementary Fig. 7a). However, only *VIM* and *KRT5* exhibited high gene expression levels in epithelial cells, owing to the limited sample amount (Supplementary Fig. S8b). To clarify the differential expression patterns, we entered these genes into a well-established database, UALCAN [32], and compared their expression levels in 114 normal samples and 1097 breast cancer samples. The expression levels of *VIM, KRT5*, and *KRT17* in normal samples were significantly higher than those in breast cancer samples (Fig. 6e). Conversely, the expression levels of *KRT8, KRT18*, and *FOXA1* showed the opposite trends. This is consistent with the results obtained from peak enrichment, thus providing compelling evidence that the predicted tumor cells and normal cells in the TME are precise and reasonable, and demonstrating the suitability of HyGAnno for researchers who wish to annotate scATAC-seq derived from tumor tissues and explore the mechanisms of the carcinogenesis from an epigenomic perspective.

## 3 Discussion

scATAC-seq, one of the most important sequencing protocols, is used to study genomewide open chromatin regions at the epigenetic level. Together with scRNA-seq at transcriptomic level, scATAC-seq offers the opportunity to unveil the intricate regulatory mechanisms across diverse cellular phenotypes [21, 33]. As a prerequisite for single-cell multi-omics analysis, cell type annotation has been well explored in scRNA-seq data, while the same problem in scATAC-seq data has received less attention. This gap limits our understanding of the roles of CREs in determining cell type-specific functionalities, such as promoter-distal open chromatin regions, which contain enhancers that regulate cell type-specific transcription [34]. To address this, we developed a deep graph-based model, HyGAnno, that transfers cell label information from scRNA-seq data to cells assayed by means of scATAC sequencing. In contrast to other cell type annotation methods that train model with only gene-level features, HyGAnno innovatively constructs advanced graph neural networks with both the peak-level and gene-level features, thus exploring the possibility of modeling transcription processes with the assistance of multi-omics data.

According to the experimental results, HyGAnno outperformed most existing cell annotation tools, especially those that overlooked the promoter-distal accessible peak information during the model training process. The interpretable cell embedding of HyGAnno provided new insights into traditional trajectory analysis, and the continuous pseudotime may reveal the key genes dynamically expressed in a specific cell differentiation process. As a label transferring method, HyGAnno relies on the cell label information in the scRNA-seq reference data, which is manually or computationally annotated based on expression similarities to some marker genes. However, spillage from cell-free ambient RNA or barcode swapping events cause background noise in droplet-based scRNA-seq experiments, thus reducing the significance of marker gene detection [35] and leading to noisy cell annotation in the reference data. To reduce the effect of these noisy cell labels, HyGAnno reconstructs a new graph to retain the backbone of the inter-modality and intra-modality cell connections in the initial graphs. Reconstruction loss can reduce the weight of the classification loss affected by noisy labels. Meanwhile, the reconstructed graph can also amplify the abnormal cell-cell connections in the initial graphs, and hence, be leveraged to judge the prediction reliability by assigning each cell a confident or an ambiguous description. When ambiguous predictions appear in a boundary or mixture consisting of multiple cell types, we suggest removing these predictions before downstream analysis. However, when ambiguous predictions are enriched in entire cell clusters, we recommend that users change other reference data or note the potential novel cell types. We also validated HyGAnno facility in TMEs. HyGAnno successfully distinguished between tumor and normal cells, thereby providing insights into the mechanisms of the carcinogenesis at epigenomic perspective. Finally, with the growing repository of high-quality scRNA-seq datasets, using scRNA-seq references that correspond to the tissue with the scATAC-seq target becomes more practical [36]. This development provided a critical foundation for HyGAnno to effectively transfer cell labels from scRNA-seq data to scATAC-seq data. We have also planned some strategies for further development of HyGAnno. First, HyGAnno can be adapted to paired experimental protocols, such as SNARE-seq [1] and SHARE-seq [2], which allow the simultaneous measurement of gene expression and chromatin accessibility in the same cell. By constructing multiple hybrid graphs and using ATAC-anchor cells as a bridge, HyGAnno can reinforce information-sharing among modalities, relieving low-throughput and sparse data [37, 38] from these joint profiling technologies. Second, as the proper selection of reference data is of vital importance for the annotation result, HyGAnno should evaluate the reference data based on the distribution of ambiguous predictions, there by advising users on the necessity of additional RNA-seq references.

## 4 Conclusions

In this study, we have showcased the capabilities of HyGAnno, a parallel graph neural network designed to transfer cell label information from scRNA-seq to scATAC-seq. Beyond just focusing on gene expression (activity) patterns across different cell types, HyGAnno places greater emphasis on understanding how genome-wide chromatin accessibility features contribute to cellular heterogeneity, which is achieved by training the model using the raw data of open chromatin regions. HyGAnno showed a significant improvement in the prediction accuracy and provided interpretable cell embeddings. Taking advantage of generative part in HyGAnno model, an informative reference-target cell graph was reconstructed, which could amplify abnormal cell-cell connections and describe the cell type prediction reliability. In addition, HyGAnno successfully distinguished neoplastic and healthy cells in the tumor microenvironment, showing its high annotation adaptability in complex tissues. We believe that HyGAnno, supported by advanced deep-learning models and graph analysis, has the potential to enhances our understanding of the role of genome-wide accessible peaks in cell differentiation and development.

## 5 Methods

### 5.1 Graph construction

After applying Seurat [8] and Signac [13] for quality control, a gene expression matrix 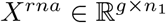 from scRNA-seq data and a peak matrix 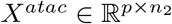 from scATAC-seq data were generated, where *g*(= 2000) is the number of highly variable genes, *p* is the number of top accessible peaks, and *n*_1_ and *n*_2_ are the cell numbers. *X*^*atac*^ was used to build a gene activity matrix (GAM) 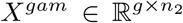, which represents the degree of accessibility in gene body regions of highly variable genes [12, 13]. HyGAnno built three undirected graphs: G^*rna*^, a cell-clustering graph of scRNA-seq; *G*^*H*^, a hybrid graph connecting *G*^*rna*^ and ATAC anchor cells; and *G*^*atac*^, a cell-clustering graph of scATAC-seq. *G*^*rna*^ was built using shared nearest neighbors coupled with the principal component analysis (PCA) with *X*^*rna*^ (nPCs=30). To detect ATAC anchor cells, we standardized *X*^*rna*^ and *X*^*atac*^ and used canonical correlation analysis [39] measures the similarity between RNA cells and ATAC cells [10]. The ATAC cells having higher similarities to the shared nearest neighbors of RNA cells become anchor cells positioned in *G* ^*rna*^, yielding 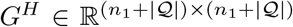 and the corresponding feature matrix 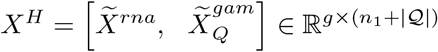, where 𝒬 is the set of ATAC anchor cells; and 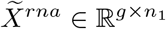 and 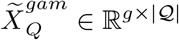 are the standardized feature matrices of *X*^*rna*^ and 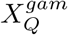 containing only ATAC anchor cells, respectively (Supplementary Table 1). *G*^*atac*^ was created by applying a shared nearest neighbor strategy on the LSI latent space. The node feature of *G*^*atac*^ is the term frequency-inverse document frequency (TF-IDF) normalized peak matrix 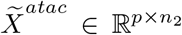. The input sets for training the neural network are summarized as the graph set 𝔾 = {*G*^*H*^, *G*^*atac*^} and feature set 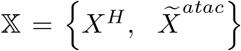. Details of the graph construction and selections of anchor cell numbers are provided in Supplementary Notes 1 and 2, respectively.

### 5.2 Parallel variational graph auto-encoders

To transfer the cell labels from the hybrid graph to an ATAC graph, a parallel variational graph auto-encoder (VGAE) [40] framework embeds graphs in 𝔾 together with 𝕏 into a k-dimensional space, as follows:

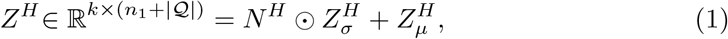

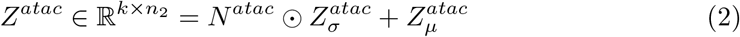

where 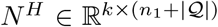 and 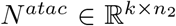 are standard normal random matrices; *A* ⊙ *B* represents the Hadamard product of two matrices; 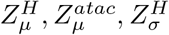 and 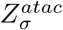 are calculated using a two-layer graph convolutional network (GCN) [41]. Finally, the loss function that regulates the latent variables *Z*^*H*^ and *Z*^*atac*^ in the two VGAEs is optimized as follows:

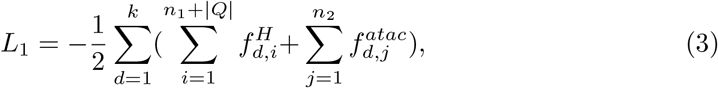

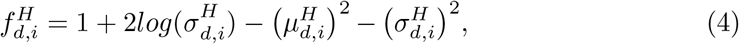

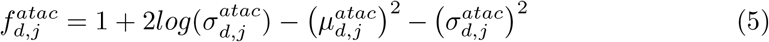

where k is the dimension number of the embedding space, 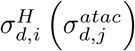 and 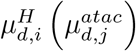 are the element in d-th row and *i*−*th*(*j*−*th*) column of the 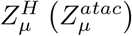 and 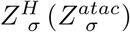. Complete descriptions are provided in Supplementary Note 3.

### 5.3 Label transferring for ATAC non-anchor cells

After graph embedding, label knowledge is propagated from the RNA cells to the ATAC anchor cells. However, non-anchor cells in the ATAC graph still lack cell labels. Therefore, we designed an alignment strategy inspired by the concept of graph contrastive learning [42, 43] by treating the ATAC anchor cells viewed by two graphs as positive node pairs. More specifically, we suspected that the same unlabeled ATAC anchor cells in different graphs have the same representation, which yields the following loss function:

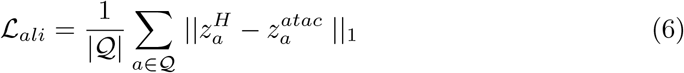

where 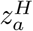 and 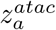 are the feature vectors of the ATAC anchor cell a of *Z*^*H*^ and *Z*^*atac*^, respectively, and || * ||_1_ is the Euclidean norm. Therefore, the networks can be collaboratively trained by using not only the gene-level information in *G*^*H*^ but also the peak-level information in *G*^*atac*^.

### 5.4 Graph integration and reconstruction

To deal with the high sparsity in *G*^*H*^ and *G*^*atac*^, we combined them to one graph 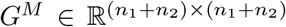 and reconstructed a new graph 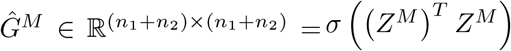, where *σ*(·) is the logistic sigmoid function, and *Z*^M^ is the concatenation of *Z*^*H*^ and 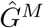 was supposed to be more informative than *G*^*M*^, as it better illustrated the correlation between RNA cells and ATAC cells (Supplementary Note 4). The graph reconstruction loss can be optimized using the binary cross entropy:

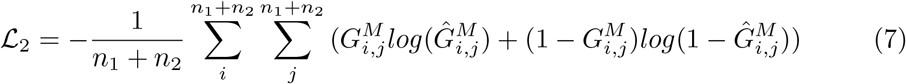

### 5.5 Loss functions in HyGAnno

We applied softmax activation on *Z*^*M*^ to compute the label indicator matrix 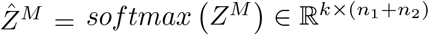. The prediction of the reference cells was then optimized using cross entropy as follows:

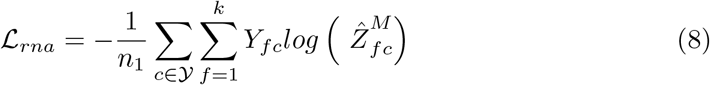

where 𝒴 is the reference cell set, 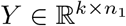 contains the true cell labels of 𝒴, and *Y*_*fc*_ = 1 indicates that the cell *c* has the *f* th label, and *Y*_*fc*_ = 0 indicates that the cell is not of this label. The final predicted label for target cell i is determined as the *f* th label when 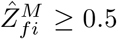. Along with other functions, the final loss function used in the HyGAnno framework is given as follows:

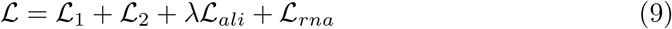

where *λ* is a parameter adjusting the strength of the alignment loss.

### 5.6 Labeling prediction certainty

The *c*th column in the label indicator matrix 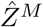 is the vector of the cell label probabilities 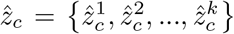, indicating that the cell *c* is likely to belong. Given an ATAC cell *c*, to describe its prediction reliability, we calculated the edge density *d* and weight *w* of cell *c* in the reconstructed reference-target cell graph, as follows:

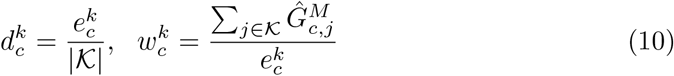

where *k* is the label indicator of the RNA cell cluster 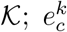 is the edge number between *c* and 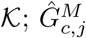 is the edge weight of the ATAC cell *c* and RNA cell *j*. When 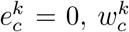 is treated as 0. These formulas yield two vectors: 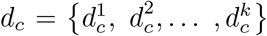 and 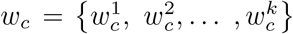. If all 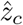, *d*_*c*_, and *w*_*c*_ achieve the highest values at the *k*th elements, the prediction of ATAC cell *c* was confident, otherwise, was ambiguous. That is, from the perspective of label probability, the label *k* with the highest 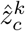 is the prediction result for the cell *c*, whereas from the perspective of graph connectivity, the highest 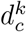 and 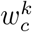 reflect that the cell *c* is densely connected to the RNA cell cluster with the label *k*. And the inconsistencies among these two perspectives were leveraged to identify ambiguous or confident predictions.

### 5.7 Input data preparation

For the RNA-referenced methods, mouse brain data (GSE126074) was downloaded from National Center for Biotechnology Information Gene Expression Omnibus (NCBI GEO) database and processed using Seurat and Signac, to create the input matrices *X*^*rna*^, *X*^*atac*^, and *X*^*gam*^. The mouse lung data was prepared from Tabula Muris [44] and the atlas of the adult mice chromatin accessibility [45] was processed by Cicero to create *X*^*gam*^. The input matrices of human PBMC and BMMC were downloaded from a previous study [21]. For the ATAC-referenced methods, GSE139369 and GSE129785 were downloaded from NCBI GEO and processed using Cicero to create *X*^*gam*^ (Supplementary Note 5). The scRNA-seq (GSE176078) and scATAC-seq (GSE198639) datasets for breast cancer data were prepared, and *X*^*gam*^ was created using Signac (Supplementary Note 6). The original sources of all data are shown in Supplementary Table 2.

### 5.8 Baseline methods

We compared HyGAnno to six cell annotation methods: Seurat (v3) [8], scJoint [9], scGCN [10], Conos [11], Cellcano [14], and EpiAnno [15]. For all methods except Cellcano and EpiAnno, we used the scRNA-seq gene expression matrix as a reference. For Cellcano, the scATAC-seq peak matrix was transformed to a gene activity matrix as a reference. For EpiAnno, we directly inputted the peak matrix as a reference and unified the open chromatin regions between the reference and target.

### 5.9 Evaluation metrics

The predictive performance was evaluated in terms of Accuracy (ACC), Normalized Mutual Information (NMI) [46], and F1 score. The F1 score was also weighted by the size of cell types in each dataset. The silhouette width (SW) score [7, 47] quantifies the capacity of a method to generate informative cell embeddings with ground truth cell types (Supplementary Note 8).

### 5.10 Downstream analyses

#### 5.10.1 TF motif enrichment

The chromVAR R package [16] version 1.18.0 was used to calculate the motif enrichment of TFs in mouse lung data. UCSC version mm9 was used as the genome reference to detect motifs, which were then used as the input for JASPAR2022, to map these motifs to TFs under experimental validation.

#### 5.10.2 Detection of cell type-specific peaks

To elaborate the relationship between peak-level and latent features, we simplified our model into one-layer neural network. After the model has been fully trained (ACC=0.90, NMI=0.80), we extracted the hidden parameter matrix from the ATAC channel. For each cell type-specific latent features, we selected the top 500 peaks according to their feature values as the cell type-specific peaks, where an overlapping threshold of 200 bases is applied to map promoter-distal peaks to cell type-specific enhancers from the scEnhancer database.

#### 5.10.3 Cell embedding generation and visualization

For PCA, the first 30 principal components of the gene activity matrix were used for cell embeddings. For LSI, Singular Value Decomposition (SVD) was applied to the term frequency-inverse document frequency (TF-IDF)-coded peak matrix, with the second to thirtieth SVD components serving as cell embeddings. For HyGAnno, scJoint, and scGCN, the latent spaces in the neural networks were treated as cell embeddings (Supplementary Note 7). All cell embeddings were visualized using UMAP, with the fixed parameters *n-neighbors=50* and *min-dist=0*.

#### 5.10.4 Trajectory inference based on cell embedding space

The destiny R package [48] version 3.10.0 was used to infer the cell trajectories. We applied the inner functions using cell embedding as the input and calculated the diffusion pseudotime with all default parameters. The first two diffusion components were used for trajectory visualization.

#### 5.10.5 CNV calculation for scATAC-seq data

We inferred the CNV signals by running the scripts provided in a previous work [29]. After merging the raw scATAC-seq fragment files from different patients, we separated the chromosomes into 10-Mb windows and returned the average log2(fold-change) of fragment count falling into each window against the 100 nearest neighbors, based on GC content. Regions with a higher log2(fold-change) were considered candidates for amplification. To clearly illustrate the CNV pattern between tumor cells and normal cells in breast cancer, we subtracted the mean value of each window from the CNV scores of TME cells and obtained the normalized CNV scores [27].

## 6 Availability of data and materials

All data used in this study were obtained from public resources and can be downloaded from original publications as mentioned in section 5.7. The codes and tutorials are freely available at https://github.com/WeihangZJP/HyGAnno under a MIT license.

## 9 Authors’ contributions

W.H.Z. contributed to the conception, design, analysis, and interpretation of the data.

C.Y. interpreted the biological explanations. B.W.L. tested and debugged the computational code. W.H.Z. drafted the manuscript. W.H.Z, S.J.P., M.L. and K.N. critically revised the manuscript. K. N. supervised the study.

## 10 Acknowledgements

Computational resources were provided by the supercomputer system SHIROKANE at the Human Genome Center, Institute of Medical Science, University of Tokyo.

