## Supplementary_Notes_Figures_Tables for "HyGAnno: Hybrid graph neural network-based cell type annotation for single-cell ATAC sequencing data"

**Supplementary Note 1: Details in graph construction**

For hybrid graph, it contains two kinds of edges: edges between RNA cells themselves and edges between RNA anchor cells and ATAC anchor cells. For the first kind of edges, we applied principal component analysis (PCA) to project the gene feature space of $\mathrm{X}^{rna}$ to a low dimensional space, which is consisted of top 30 PCs. Then, if any of the two RNA cells in this latent space are the shared nearest neighbors (SNNs), an edge is assigned to them. For the second kind of edges, we measured the similarity between cells in different modalities by transforming the peak matrix $\mathrm{X}^{atac}$ to the gene activity matrix (GAM) $\mathrm{X}^{gam} \in \mathbb{R}^{g \times n_2}$, which represents the degree of accessible gene body regions scored by the total number of mapped scATAC-seq reads[1]. Depending on different datasets, the GAMs may be directly provided from original works or be generated through other functions such as Signac and Cicero. After taking intersected genes of $\mathrm{X}^{rna}$ and $\mathrm{X}^{gam}$, we performed a similar standardization transform mentioned in scGCN on these two matrices and conducted canonical correlation analysis (CCA) to project them into a low dimension space, where the projected data achieves the highest correlation coefficient. Any RNA and ATAC cells clustered together by SNN are assigned as RNA anchor cells and ATAC anchor cells for the corresponding clusters. Finally, we got the hybrid graph $\mathrm{G}^{H} \in \mathbb{R}^{(n_1+|Q|)\times(n_1+|Q|)}$ and the feature matrix $\mathrm{X}^{H} = \left[\widetilde{\mathrm{X}}^{rna},\ \widetilde{\mathrm{X}}_{Q}^{gam}\right] \in \mathbb{R}^{g\times(\mathrm{n}_1+|Q|)}$, where $G_{ij}^{H} = G_{ji}^{H} = 1$ stands the edge of the cell *i* and *j*; $Q$ is the set of ATAC anchor cell; $\widetilde{\mathrm{X}}^{rna} \in \mathbb{R}^{g\times n_1}$ and $\widetilde{\mathrm{X}}_{Q}^{gam} \in \mathbb{R}^{g\times|Q|}$ are the whole and subset of the standardized feature matrix of $\mathrm{X}^{rna}$ and $\mathrm{X}^{gam}$, respectively.

For ATAC graph, a term frequency-inverse document frequency (TF-IDF) transformation followed by singular value decomposition (SVD) is performed to $\mathrm{X}^{atac}$, also known as the latent semantic indexing (LSI). We removed the first LSI component as it might capture the technical variation rather than biological variation. Similarly, we conducted SNN strategy on the LSI latent space with remained components, obtaining ATAC graph $\mathrm{G}^{atac} \in \mathbb{R}^{n_2\times n_2}$. To train HyGAnno with peak-level information, instead of frequently use the gene-level matrix of $\mathrm{X}^{gam}$, the peak-level matrix of $\widetilde{\mathrm{X}}^{atac} \in \mathbb{R}^{p\times n_2}$ is treated as the node feature of the ATAC graph, where $\widetilde{\mathrm{X}}^{atac}$ is the TF-IDF normalized matrix of $\mathrm{X}^{atac}$.

**Supplementary Note 2: Number of the anchor cells**

As a bridge of the hybrid graph and ATAC graph, the number of anchor cells can affect the cell annotation performance a lot. We found a larger shared nearest neighbor (SNN) number (around 20 in this research) in the canonical correlation analysis (CCA) can strongly improve the precision by detecting a higher number of anchor cells. This causes the anchor cells take over most of the cells in reference and target data as recorded in Supplementary Table 1. This is reasonable in our approach since our model minimizes the distance between the ATAC anchor cells viewed by two different graphs instead of the distance between the RNA anchor cells and ATAC anchor cells in the same hybrid graph. Thus, more anchor cells can broaden the bridge between hybrid graph and ATAC graph, passing more peak-level information to the hybrid graph and improve the cell annotation results. Moreover, although a larger SNN number lead to the generation of a much more complex hybrid graph, which may introduce noise edges, the graph encoding layer in our framework can extract the most informative edges and remove the useless edges in the graph reconstruction step.

**Supplementary Note 3: Graph embedding by VGAE auto-encoders**

Regarding each of the hybrid graph and the ATAC graph, we embed it into a $k$-dimensional space. In each space, to satisfy the loss function striction of the VGAE, the cell embedding should be hypothetically sampled from a specific $k$-multivariate normal distribution with independent variables:

$$q(\mathrm{Z}^H|\mathrm{X}^H, \mathrm{G}^H) = \prod_{i=1}^{k} \mathcal{N}\left(z_i^H \middle| \mu_i^H, diag(\sigma_i^H)\right),$$

$$q\left(\mathrm{Z}^{atac}\middle|\widetilde{\mathrm{X}}^{atac}, \mathrm{G}^{atac}\right) = \prod_{j=1}^{k} \mathcal{N}\left(z_j^{atac} \middle| \mu_j^{atac}, diag(\sigma_j^{atac})\right)$$

where $\mathrm{Z}^H \in \mathbb{R}^{k\times(n_1+|\mathcal{Q}|)}$ and $\mathrm{Z}^{atac} \in \mathbb{R}^{k\times n_2}$ are the embeddings of graph $\mathrm{G}^H$ $and$ $\mathrm{G}^{atac}$, respectively; $k$ is the number of the cell types in reference data; $\mu_i^H$ and $\sigma_i^H$, which are k-dimensional vectors, represent the mean and covariance of the multivariate normal distribution, respectively, so are the $\mu_j^{atac}$ and $\sigma_j^{atac}$. Then, to obtain the $\mu^H$ and $\sigma^H$ in practice, two-layer graph convolutional networks (GCNs) is conducted on hybrid graph:

$$\mathrm{Z}_\mu^H = (GCN_\mu^H(\mathrm{X}^H, \mathrm{G}^H))^T = (\widetilde{\mathrm{G}}^H ReLU(\widetilde{\mathrm{G}}^H (\mathrm{X}^H)^T \mathrm{W}_0^H)\mathrm{W}_\mu^H)^T,$$

$$\mathrm{Z}_\sigma^H = (GCN_\sigma^H(\mathrm{X}^H, \mathrm{G}^H))^T = (\widetilde{\mathrm{G}}^H ReLU(\widetilde{\mathrm{G}}^H (\mathrm{X}^H)^T \mathrm{W}_0^H)\mathrm{W}_\sigma^H)^T$$

where $\mathrm{Z}_\mu^H, \mathrm{Z}_\sigma^H \in \mathbb{R}^{k\times(n_1+|\mathcal{Q}|)}$ share the same first-layer parameter matrix of $\mathrm{W}_0^H$ but different second-layer parameter matrices of $\mathrm{W}_\mu^H$ and $\mathrm{W}_\sigma^H$; $ReLU(\cdot) = max(0\ ,\ \cdot)$ is the activation function; $\widetilde{\mathrm{G}}^H = \mathrm{D}^{-\frac{1}{2}}\mathrm{G}^H\mathrm{D}^{-\frac{1}{2}}$ is the symmetrically normalized adjacency matrix aiming to effective training, where D and I are the diagonal degree matrix of $\mathrm{G}^H$ and the identity matrix, respectively. Similarly, $\mathrm{Z}_\mu^{atac}$, $\mathrm{Z}_\sigma^{atac} \in \mathbb{R}^{k\times n_2}$ can be calculated in the same way. Then, the cell embeddings of the hybrid graph and the ATAC graph can be obtained from:

$$\mathrm{Z}^H = N^H \odot \mathrm{Z}_\sigma^H + \mathrm{Z}_\mu^H,$$

$$\mathrm{Z}^{atac} = N^{atac} \odot \mathrm{Z}_\sigma^{atac} + \mathrm{Z}_\mu^{atac}$$

where $N^H \in \mathbb{R}^{k\times(n_1+|\mathcal{Q}|)}$ and $N^{atac} \in \mathbb{R}^{k\times n_2}$ are random matrices generated form standard normal distributions; A $\odot$ B represents the Hadamard product of two matrices. Finally, the loss function which regulates the latent variable $\mathrm{Z}^H$ and $\mathrm{Z}^{atac}$ from Gaussian distribution is optimized as:

$$\mathcal{L}_1 = -(KL(q(\mathrm{Z}^H|\mathrm{X}^H, G^H)|, p(\mathrm{Z}^H)) + KL\left(q\left(\mathrm{Z}^{atac}\middle|\widetilde{\mathrm{X}}^{atac}, \mathrm{G}^{atac}\right)\middle|, p(\mathrm{Z}^{atac})\right))$$

where $KL\big(q(\cdot)\big|, p(\cdot)\big)$ is the Kullback-Leibler (KL) divergence between distribution $q(\cdot)$ and $p(\cdot)$; As Gaussian priors, $p(\mathrm{Z}^{H}) = \prod_i \mathcal{N}(z_i|0, I)$ and $p(\mathrm{Z}^{atac}) = \prod_j \mathcal{N}\big(z_j\big|0, I\big)$. According to the solution B in VAE[2], since both prior $p(\mathrm{Z}^{H}) = \prod_i \mathcal{N}(z_i|0, I)$ and posterior approximation $q(\mathrm{Z}^{H}|\mathrm{X}^{H}, \mathrm{G}^{H})$ are Gaussian, $KL\big(q(\mathrm{Z}^{H}|\mathrm{X}^{H}, G^{H})\big|, p(\mathrm{Z}^{H})\big)$ can be computed as:

$$KL\big(q(\mathrm{Z}^{H}|\mathrm{X}^{H}, G^{H})\big|, p(\mathrm{Z}^{H})\big) = -\frac{1}{2}\sum_{d=1}^{k}\sum_{i=1}^{n_1+|Q|}(1 + 2log(\sigma_{d,i}^{H}) - (\mu_{d,i}^{H})^2 - (\sigma_{d,i}^{H})^2)$$

where k is the dimension number of the embedding space, $\sigma_{d,i}^{H}$ and $\mu_{d,i}^{H}$ are the element in d-th row and i-th column of $\mathrm{Z}_{\mu}^{H}$ $and$ $\mathrm{Z}_{\sigma}^{H}$, respectively.

Similarly,

$$KL\left(q\big(\mathrm{Z}^{atac}\big|\tilde{\mathrm{X}}^{atac}, \mathrm{G}^{atac}\big)\Big|, p(\mathrm{Z}^{atac})\right) = -\frac{1}{2}\sum_{d=1}^{k}\sum_{j=1}^{n_2}(1 + 2log(\sigma_{d,j}^{atac}) - (\mu_{d,j}^{atac})^2 - (\sigma_{d,j}^{atac})^2)$$

where k is the dimension number of the embedding space, $\sigma_{d,j}^{atac}$ and $\mu_{d,j}^{atac}$ are the element in d-th row and i-th column of $\mathrm{Z}_{\mu}^{atac}$ $and$ $\mathrm{Z}_{\sigma}^{atac}$, respectively. Thus, loss $\mathcal{L}_1$ can be rewritten as:

$$\mathcal{L}_1 = -\frac{1}{2}\sum_{d=1}^{k}(\sum_{i=1}^{n_1+|Q|}(1 + 2log(\sigma_{d,i}^{H}) - (\mu_{d,i}^{H})^2 - (\sigma_{d,i}^{H})^2) + \sum_{j=1}^{n_2}(1 + 2log(\sigma_{d,j}^{atac}) - (\mu_{d,j}^{atac})^2 - (\sigma_{d,j}^{atac})^2))$$

where k is the dimension number of the embedding space, $\sigma_{d,i}^{H}$ $(\sigma_{d,j}^{atac})$ and $\mu_{d,i}^{H}$ $(\mu_{d,j}^{atac})$ are the element in d-th row and i-th (j-th) column of $\mathrm{Z}_{\mu}^{H}$ $(\mathrm{Z}_{\mu}^{atac})$ $and$ $\mathrm{Z}_{\sigma}^{H}$ $(\ \mathrm{Z}_{\sigma}^{atac})$, respectively.

**Supplementary Note 4: Differences between the initial and reconstructed RNA-ATAC graph**

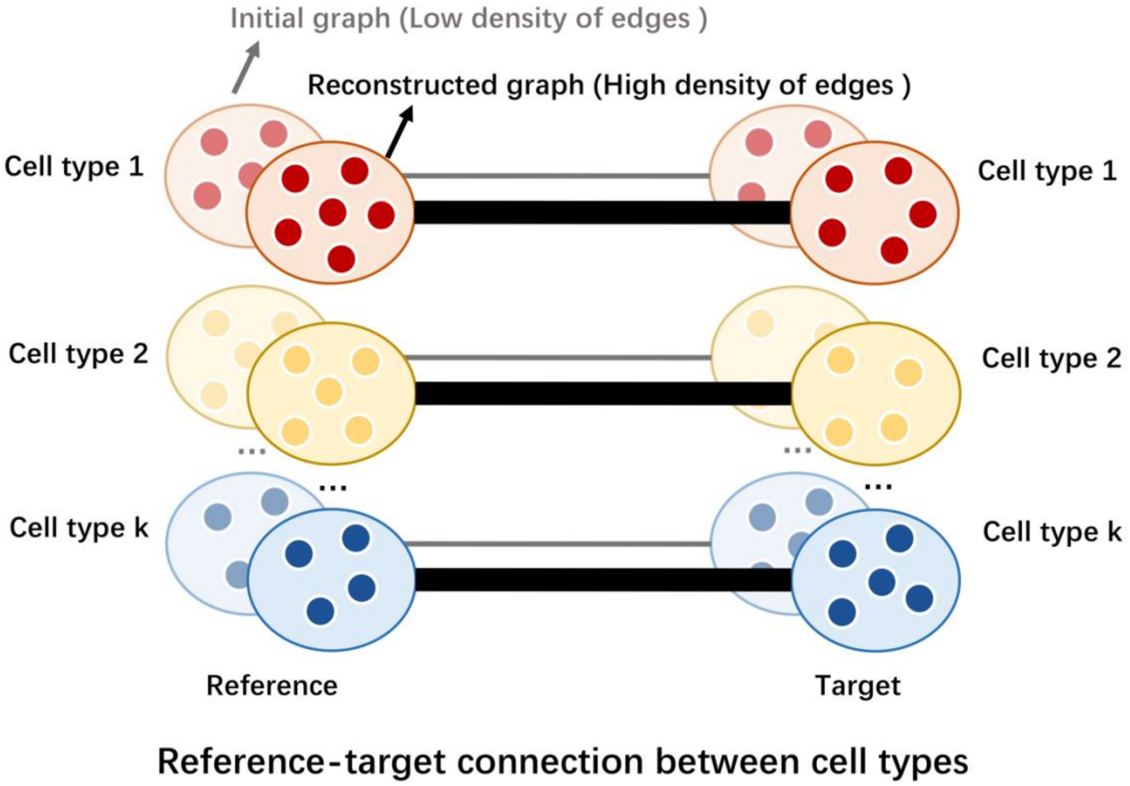

Firstly, we elaborated the role of the reconstructed graph in our model. Since our model is based on the Variational Graph Autoencoder (VGAE) which is the graph version of the Variational Autoencoder (VAE) model[2], it is crucial to understand the workings of VAE. The VAE model embeds the original data space into a latent space and assumes that different categories are sampled from a multivariate gaussian distribution. It then uses the latent space to reconstruct the original space and minimizes the difference between the initial one and reconstructed one. This ensures that the trained latent space contains most of the information of the original space. Following a similar approach, VGAE takes the initial graph as input, projects it into the latent space and reconstructs a new graph. According to previous research[3], the reconstructed graph produced by VGAE is typically used to perform link prediction tasks. This is because, compared to the initial graph, the reconstructed graph possesses a higher density of edges, and is therefore more informative. In the context of HyGAnno, this rich, detailed information allows for more accurate and robust cell annotation.

Then, in our research, the edges of reconstructed RNA-ATAC graph $G$ are supposed to be highly linked between the cells in reference and target. To prove the reconstructed graph has higher density of edges than that of the initial one, we calculated the density of edges (DLE) between cell type *i* in reference and cell type *j* in target by the following metric:

$$DLE_{i,j} = \frac{\sum \hat{G}_{c_i^{ref}, c_j^{tar}}}{N_i^{ref} N_j^{tar}}$$

where $N_i^{ref}$ and $N_j^{tar}$ are the size of cell type *i* in reference and cell type *j* in target, respectively; $\hat{G}_{c_i^{ref}, c_j^{tar}}$ is the edge weight between two cells with the cell type *i* of cell and *j*. The DLE values of the initial and reconstructed graph are calculated and visualized to demonstrate the higher density of edges in reconstructed graph (Supplementary Fig. 4).

### Supplementary Note 5: Details of data preparation for benchmarking

***Datasets for RNA-referenced methods.*** We downloaded the processed SNARE-seq data of the mouse brain from the NCBI GEO (GSE126074) and analyzed cell clusters using Seurat. This provided 14 cell clusters, and we annotated the clusters based on the marker gene expression and functions mentioned in the original publication[4]. To prepare the chromatin accessibility matrix, we remained the ATAC-seq peaks included in larger than 10 cells in the SNARE-seq data. To create the gene activity matrix, we run *GeneActivity* function in Signac to estimate reads mapped within the 2-Kb upstream and gene-body regions. The intersection of highly variable genes from the gene expression matrix and gene activity matrix are used as features. As a result, we created the gene expression matrix ($\mathrm{X}^{rna}$) of 1,305-genes-by-8,055-cells, the chromatin accessibility matrix ($\mathrm{X}^{atac}$) of 64,064-peaks-by-8,055-cells, and the gene activity matrix ($\mathrm{X}^{gam}$) of 1,305-genes-by-8,055-cells. For the mouse lung dataset, the scRNA-seq and scATAC-seq data are downloaded from the database of *Tabula Muris*[5] and the atlas of the adult mice chromatin accessibility[6], respectively. The gene activity matrix calculated by Cicero[7] is obtained directly from the data website same as the scATAC-seq data. As a result, we prepared $\mathrm{X}^{rna}$ of 1,822-genes-by-2,623-cells, $\mathrm{X}^{atac}$ of 51,465-peaks-by-7,499-cells, and $\mathrm{X}^{gam}$ of 1,822-genes-by-7,499-cells with 8 cell types. For human peripheral blood mononuclear cells (PBMC) datasets, we processed the scRNA-seq and scATAC-seq data with 10 cell types from GSE139369 based on a previous research[8] and obtained $\mathrm{X}^{rna}$ of 1,825-genes-by-13,345-cells, $\mathrm{X}^{atac}$ of 24,322-peaks-by-7,828-cells, and $\mathrm{X}^{gam}$ of 1,825-genes-by-7,828-cells. The gene activity matrix calculated by Cicero is downloaded together with the scATAC-seq data. For bone marrow mononuclear cells (BMMC) datasets, we downloaded the raw data from the same repository as PBMC and finally obtained $\mathrm{X}^{rna}$ of 1,788-genes-by-11,884-cells, $\mathrm{X}^{atac}$ of 21,201-peaks-by-14,753-cells, and $\mathrm{X}^{gam}$ of 1,788-genes-by-14,753-cells with 13 cell types.

***Datasets for ATAC-referenced methods.*** We used the scATAC-seq PBMC datasets from three healthy donors[8,9]: D10, D12, and $D_{rep1}$, which are 452,004-peaks-by-2,588-cells with 9 cell types, 452,004-peaks-by-3,070-cells with 9 cell types, and 127,541-peaks-by-9,060-cells with 6 cell types, respectively. Among them, D10 and D12 are downloaded from GSE139369, $D_{rep1}$ is downloaded from GSE129785. For convenience, we renamed the peak matrices of D10, and D12, and $\mathrm{D_{rep1}}$ as peak matrices 1, 2, and 3, respectively. As Cellcano requires gene-level matrices as inputs, the gene activity matrix of each peak matrix is necessary. For peak matrices 1 and 2, the corresponding gene activity matrices calculated by Cicero are already provided in the original research. The gene activity matrix of peak matrix 3 is unavailable, prompting us to generate it ourselves using the pipeline recommended by Cicero. Consequently, the gene activity matrices of peak matrices 1, 2, and 3 are named as gene activity matrices 1, 2, and 3, respectively. On the other hand, EpiAnno uses peak-level matrices as inputs, and it specifically requires that the peak regions in the reference and target are identical. To satisfy this requirement, we regenerate the target peak matrix through the *FeatureMatrix* function in the Signac package based on the peaks of the reference data and the fragment file of the target data acquired from the original work.

**Supplementary Note 6: Details of data preparation for tumor cell detection**

The scRNA-seq data used in tumor cell detection task was obtained from a previous study[10]. In this work, the authors analyzed 26 primary tumors with three broad subtypes, including luminal, HER2+, and triple negative breast cancer (TNBC). After the initial quality control steps, we ended up with a dataset consisting of 75,443 single cells. Because the neoplastic cells and normal epithelial cells often exhibit similar gene expression pattern, it can be challenging to annotate them based only on gene markers. To address this, we employed a method called scATOMIC, a pan-cancer classifier that has been trained on more than 300,000 cells[11], to estimate single cell copy number variant (CNV) profiles. With the help of scATOMIC, we identified seven cell types, including tumor cells. To accelerate the training of HyGAnno, we used a subset of the original scRNA-seq data, resulting $\mathrm{X}^{rna}$ of 1,539-genes-by-15,088-cells, with 6 healthy cell types and 1 tumor cell type.

For scATAC-seq data, we selected dataset derived from16 patients with three broad cancer subtypes, consistent with the scRNA-seq mentioned above. Due to the inaccessibility of the peak matrix and cell annotation information in the original study, we decided to process the raw data ourselves. First, we merged fragment files derived from 16 patients. Then, *CallPeaks* function and *FeatureMatrix* function in the Signac package are employed to enrich reads to peaks and generate peak matrix, respectively. While scRNA-seq data allowed for the usage of CNV profiles to differentiate between tumor cells and normal epithelial cells, there is no such direct method exists for scATAC-seq. This is due to scATAC-seq data quantifies the open accessibility level of chromatin regions, rather than gene expression like scRNA-seq. Hence, instead of high-resolution annotation, we transferred peak matrix to gene activity matrix by *GeneActivity* function of Signac package and roughly annotated cells by marker genes of normal cells, identifying six cell types except the tumor cell. Finally, $\mathrm{X}^{atac}$ of 65,526-peaks-by-11,116-cells, and $\mathrm{X}^{gam}$ of 1,539-genes-by-11,116-cells with 6 healthy cell types are obtained.

**Supplementary Note 7: Cell embedding generation methods**

Given the sparse and high-dimensional peak matrix in scATAC-seq data, directly exploring cell similarity in the original cell space is challenging. However, cell embedding space, a low-dimensional representation of the original space, can assist researchers in investigating the chromatin accessibility differences between the cell types. There are various ways to generate the cell embedding space, here we mainly showed the comparison methods used in this paper.

For cell embeddings with the latent semantic indexing (LSI) method, we used the functions available in Signac[1]. First, term frequency-inverse document frequency (TF-IDF) normalization is performed on the raw peak matrix using *RunTFIDF* function. The TF-IDF normalized peak matrix is then subjected to singular value decomposition (SVD) via the *RunSVD* function. Following the instructions of Signac, we discarded the first SVD component, as it typically captures technical variation rather than biological variation. We then used the SVD components from top 2 to 29 to compute the ASW scores.

For cell embeddings with PCA, we followed the pipeline suggested by Seurat[12] and applied PCA on the gene activity matrix with 2000 highly variable genes. The first 30 PCs are used to compute the ASW scores.

For cell embeddings with scJoint[13] and scGCN[14], we executed the scripts provided by the authors with default parameters. Since these methods employ deep neural networks, the output layers with 64 and 32 dimensions are considered as cell embeddings for scJoint and scGCN, respectively. Both of methods automatically output cell embedding files, which can be used to compute the ASW scores.

For our method, HyGAnno, similar with scJoint and scGCN, we used the output layer as the cell embedding to calculate the ASW scores. Different from the scGCN and scJoint, we set the dimension number of the output layer equal to the cell type number of reference data.

**Supplementary Note 8: Evaluation metrics**

***Evaluation of cell annotation performance***. Assuming that the ground truth of target data and the predicted cell label list by different methods are **X** and **Y**, respectively. For the general evaluation of the cell annotation performance, we used Accuracy (ACC) and Normalized Mutual Information (NMI):

$$ACC(X,Y) = \frac{Correct\ cell\ numbers}{All\ cell\ numbers} \in [0,1]$$

$$NMI = \frac{MI(\boldsymbol{X},\boldsymbol{Y})}{\sqrt{H(\boldsymbol{X}) \cdot H(\boldsymbol{Y})}} \in [0,1]$$

where $MI(\cdot,\cdot)$ is the mutual entropy between two list; $H(\cdot)$ is the entropy of a list. In this NMI formulation, we took the geometric mean to normalize the mutual information.

We also used F1 score to elaborate the prediction performance in cell type *i*:

$$F1_i = 2 \cdot \frac{Precision_i \cdot Recall_i}{Precision_i + Recall_i} \in [0,1]$$

where $Precision_i = \frac{TP_i}{TP_i+FP_i}$; $Recall_i = \frac{TP_i}{TP_i+FN_i}$; $TP_i$, $FP_i$, and $FN_i$ are the number of true positive cells, false positive cells, and false negative cells of cell type *i*, respectively. Also, we have the weighted F1 score normalized by the size of different cell types:

$$Weighted\ F1\ score = \frac{\sum_{i=1}^{k} F1_i \cdot c_i}{\sum_i^k c_i} \in [0,1]$$

where $F1_i$ and $c_i$ are the F1 score and the cell number of cell type *i*, respectively, k is the number of cell types. Similarly, we have the average F1 score:

$$Avg.F1\ score = \frac{\sum_{i=1}^{k} F1_i}{k} \in [0,1]$$

***Evaluation of cell embedding performance.*** The Silhouette Width (SW)[15] is a metric that quantifies the quality of clustering assignments. For a given cell, it calculates the average distance to other cells in its own cluster (inner-cluster distance) and compares this to the average distance to cells in the other cluster (inter-cluster distance). We calculated the SW score for each cell type *i* as:

$$SW_i = \frac{b-a}{max(a,b)} \in [-1,1]$$

where $a$ is the inner-cluster distance and $b$ is the inter-cluster distance. $ASW = \frac{1}{k}\sum_{i=1}^{k} SW_i$ is the average SW among all cell types, where *k* is the number of cell types. Higher $ASW$ means well-separate cell clusters, that cells with the same labels are close with each other; cells with distinct labels are far away from each other.

### Supplementary Figures

### Supplementary Figure 1

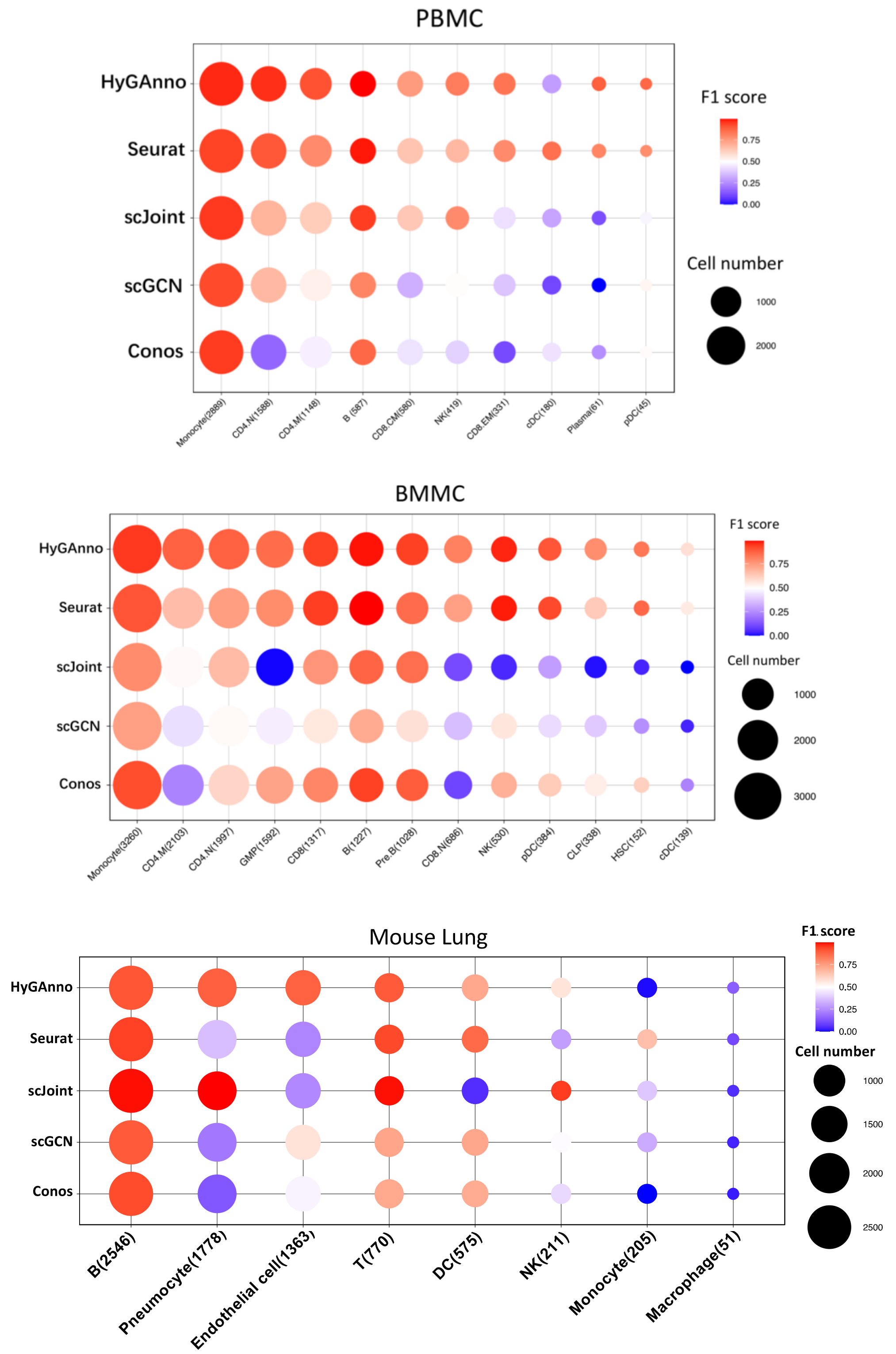

**Supplementary Figure 1**: Comparisons of F1 scores regarding each cell type in PBMC, BMMC, and mouse lung data. Number in bracket is the cell number of each cell type.

### Supplementary Figure 2

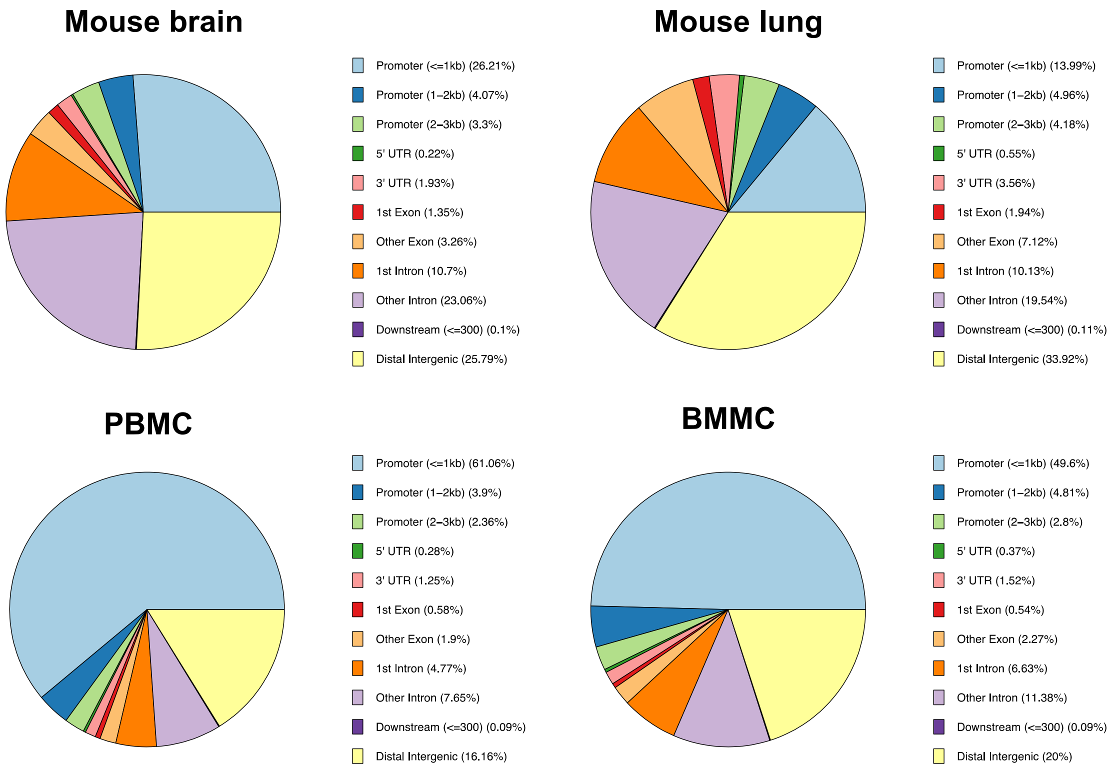

**Supplementary Figure 2**: Peak annotation for the genome-wide accessible peaks of four datasets used in HyGAnno.

**Supplementary Figure 3**

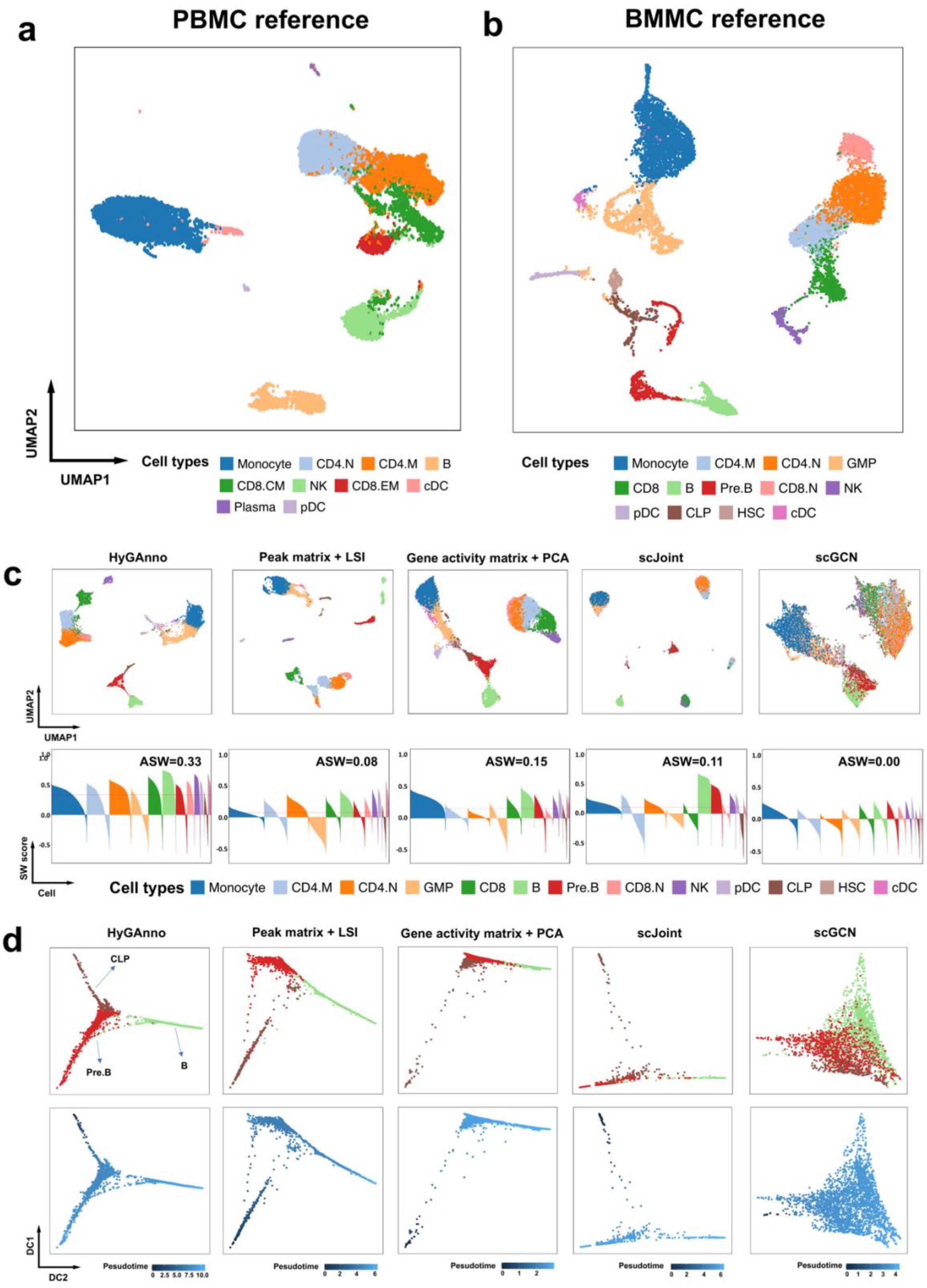

**Supplementary Figure 3**: **(a, b)** UMAP plots of cell embeddings in scRNA-seq reference data of PBMC and BMMC, respectively. Cell embeddings are obtained by applying PCA on the gene expression matrix. **(c)** UMAP plots (upper row) and the corresponding SW scores (bottom row) of BMMC cell embeddings. The cells are colored by ground truth cell types. Four T cell subtypes are surrounded by red circle. **(d)** Trajectory analysis based on the cell embeddings of CD4.N cells and CD4.N cells. The cells are colored by ground truth cell types (upper row) and pseudotime calculated by DPT (bottom row).

### Supplementary Figure 4

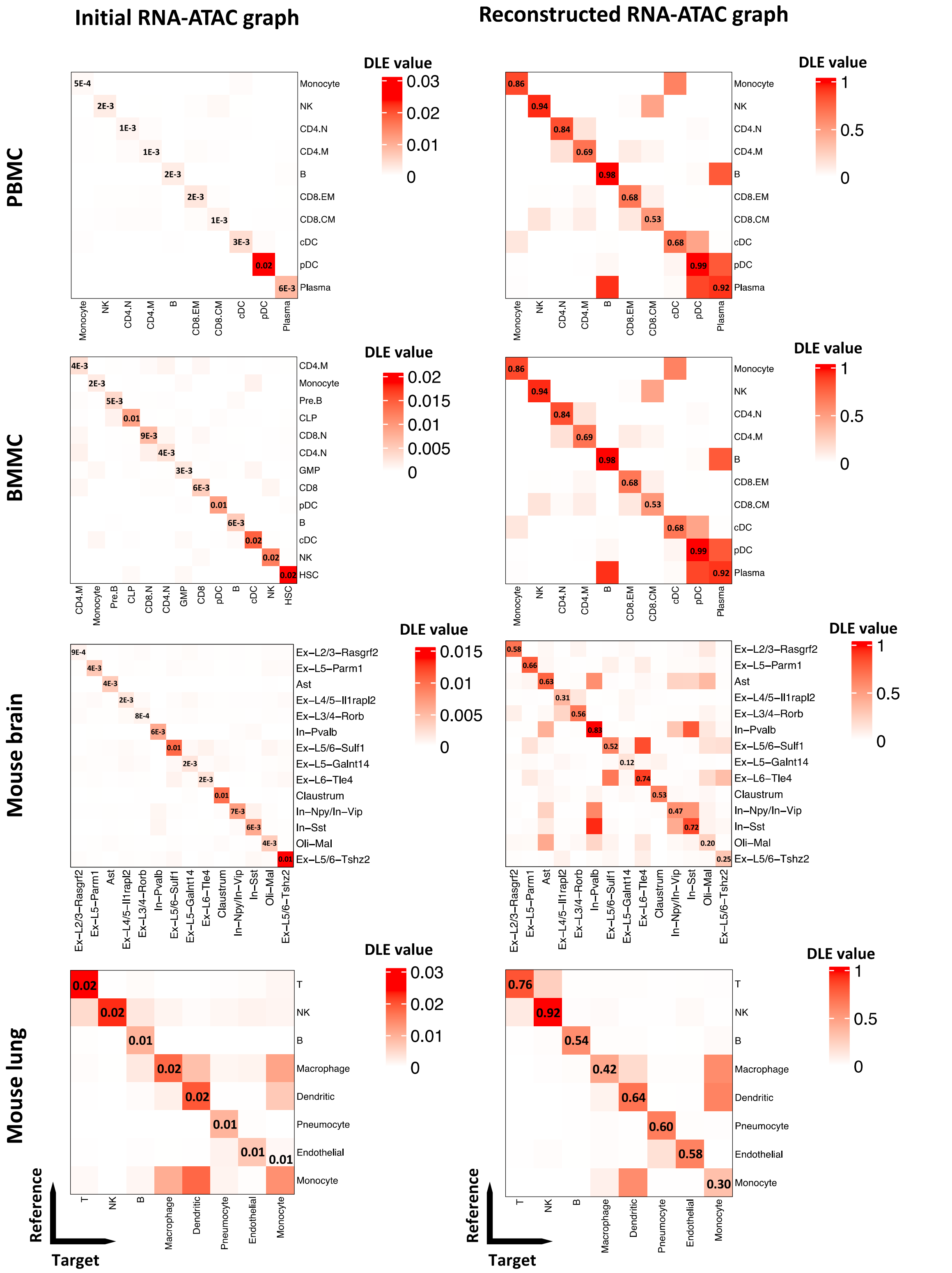

**Supplementary Figure 4**: Reconstructed RNA-ATAC graphs show higher density of edges compared with the initial one. Rows are the cells in reference data; columns are the cells in target data.

### Supplementary Figure 5

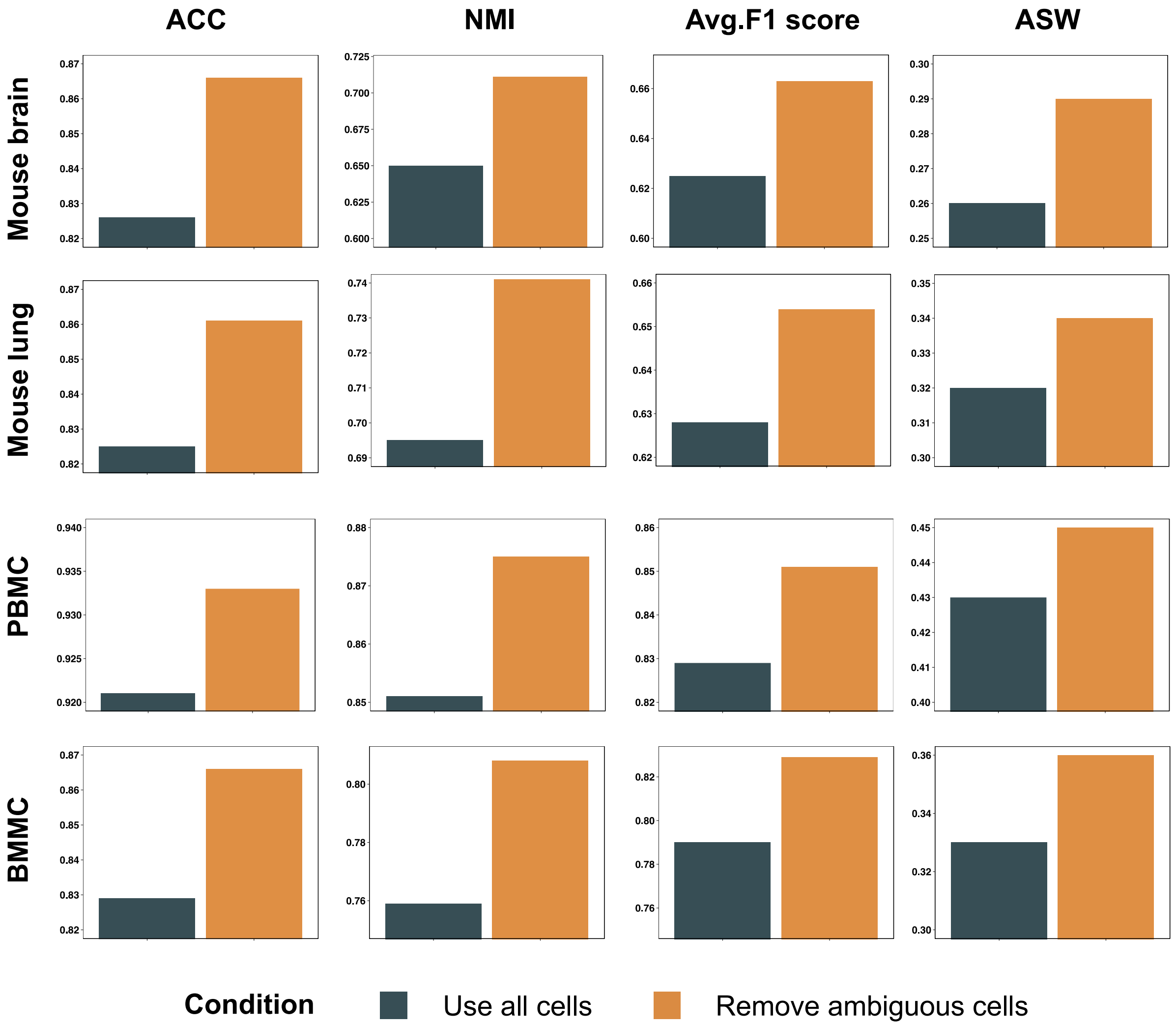

**Supplementary Figure 5**: Cell annotation performance comparisons with and without the ambiguous cells by ACC, NMI, average F1 score, and ASW.

**Supplementary Figure 6**

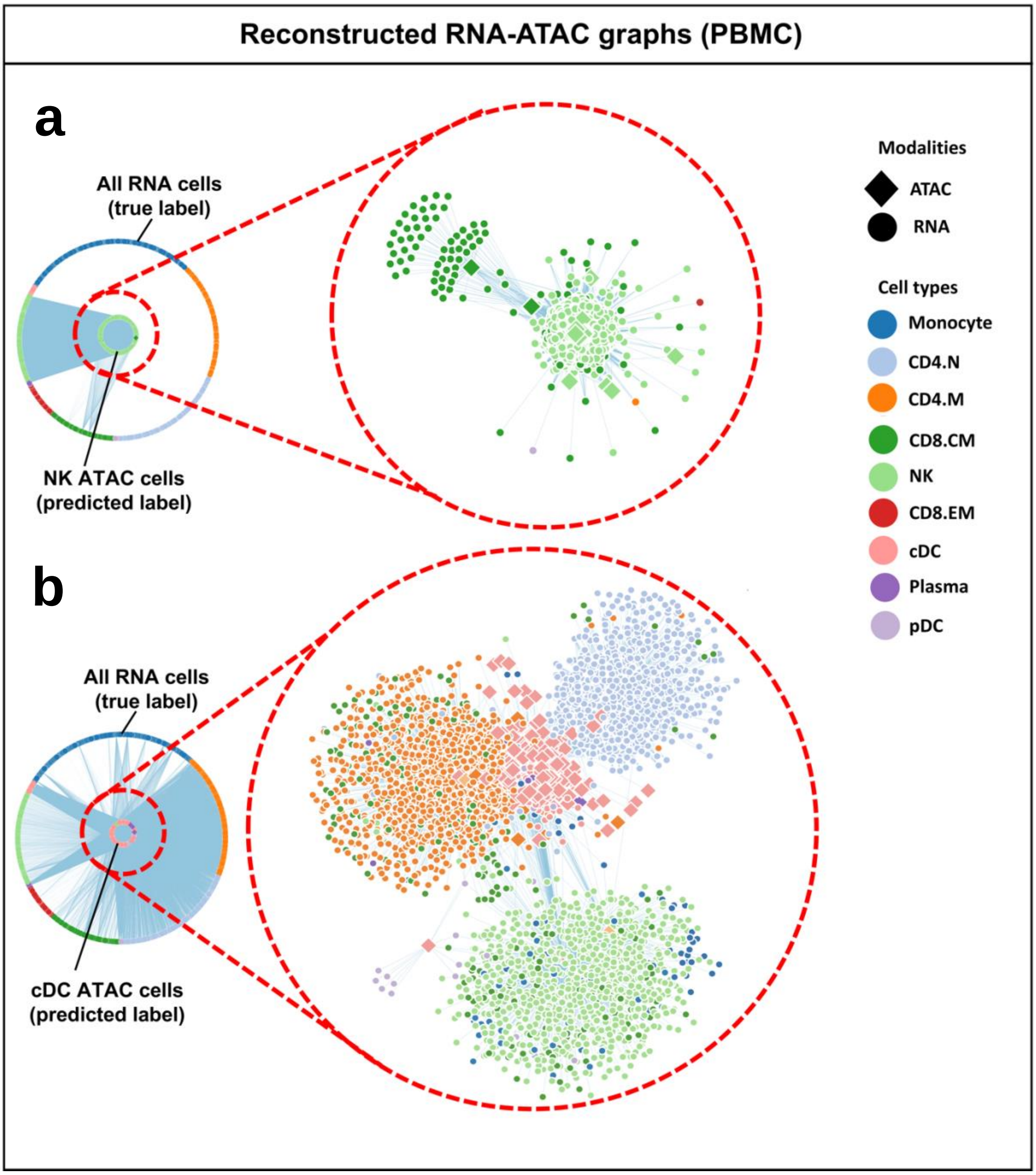

**Supplementary Figure 6**: (**a)** The reconstructed graph between confident cells and RNA clusters in PBMC data which are colored by predicted and true cell types, respectively. (**b)** The connectivity pattern between ambiguous cells and RNA clusters in PBMC data which are colored by predicted and true cell types, respectively.

### Supplementary Figure 7

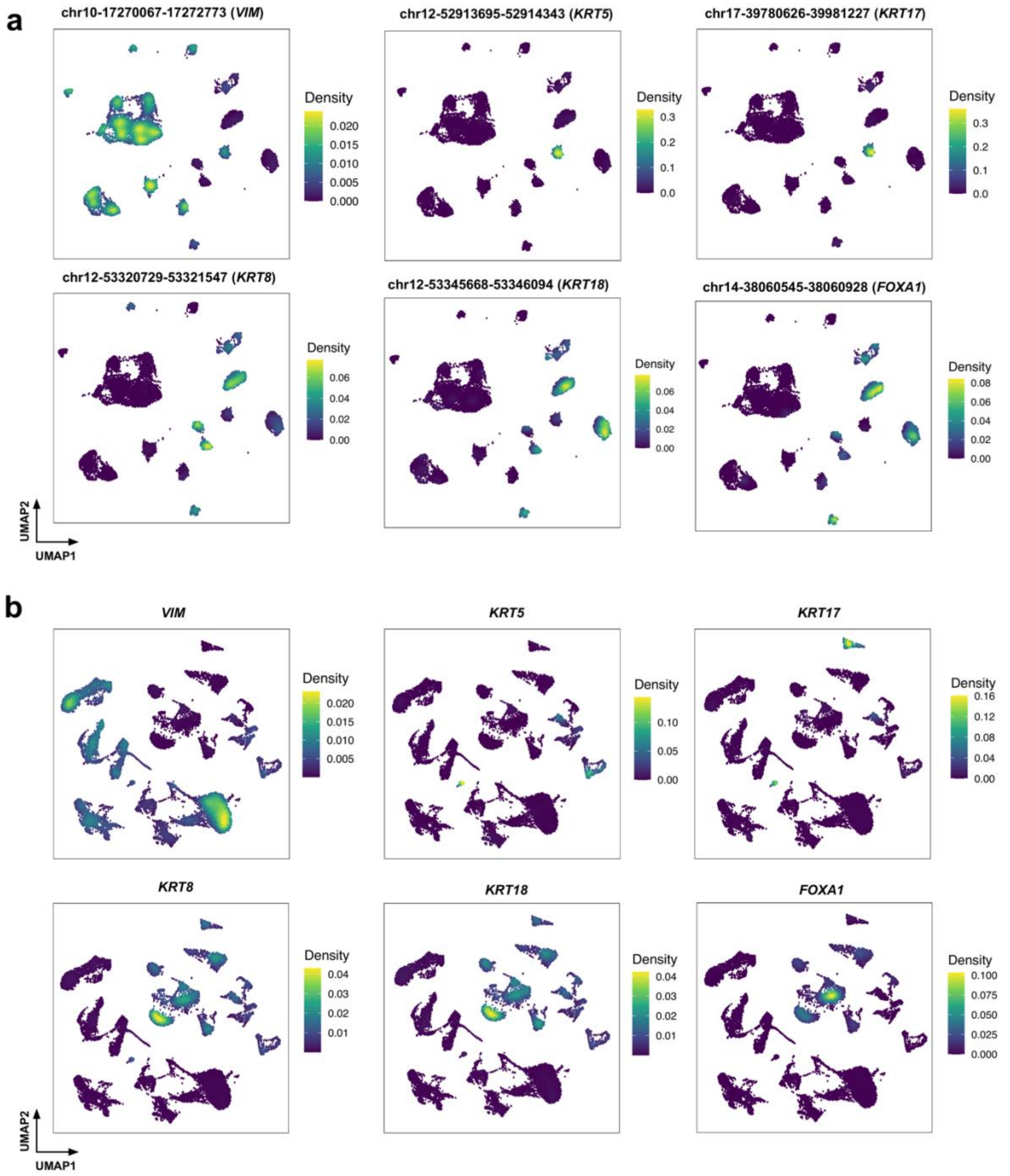

**Supplementary Figure 7:** (**a**) The accessibility level of peaks in the promotor regions of significant genes in scATAC-seq data. (**b**) The expression level of significant genes in scRNA-seq data.

### Supplementary Tables

### Supplementary Table 1: Number of anchor cells used in reference and target data

| Experiment | Node num. (RNA graph) | Edge num. (RNA graph) | Node num. (ATAC graph) | Edge num. (ATAC graph) | Anchor cell num.(RNA) | Anchor cell num.(ATAC) | Linked edge num. |
|---|---|---|---|---|---|---|---|
| Mouse brain | 8,055 | 50,028 | 8,055 | 103,724 | 7,577 | 7,349 | 22,959 |
| Mouse lung | 2,623 | 33,131 | 7,499 | 85,966 | 2,618 | 6,305 | 49,115 |
| PBMC | 13,345 | 164,947 | 7828 | 112,848 | 10,911 | 6,949 | 23,725 |
| BMMC | 11,884 | 157,846 | 14,753 | 221,550 | 11,850 | 12,725 | 122,025 |
| Breast cancer | 15,088 | 150,044 | 11,116 | 131,879 | 14,175 | 9,324 | 84,520 |

### Supplementary Table 2: Dataset sources and descriptions

| Dataset | Type | Usage | Cell num. | Gene/Peak num. | Cell type num. | Gene score strategy | Source |
|---|---|---|---|---|---|---|---|
| Mouse brain (mm10) | RNA | Reference | 8,055 | 33,160 | 14 | No need | Chen et al., 2019 |
| Mouse lung (mm10) | RNA | Reference | 2,623 | 23,433 | 8 | No need | Schaum et al., 2018 |
| PBMC (hg19) | RNA | Reference | 13,345 | 20,287 | 10 | No need | Granja et al., 2019 |
| BMMC (hg19) | RNA | Reference | 11,884 | 20,287 | 13 | No need | Granja et al., 2019 |
| Breast cancer (hg19) | RNA | Reference | 15,088 | 27,719 | 7 | No need | Wu et al., 2021 |
| PBMC Rep1 (hg19) | ATAC | Reference | 9,060 | 127,541 | 6 | Ciero | Satpathy et al., 2019 |
| Mouse brain (mm10) | ATAC | Target | 8,055 | 267,670 | 14 | Signac | Chen et al., 2019 |
| Mouse lung (mm9) | ATAC | Target | 7,499 | 436,206 | 8 | Ciero | Cusanovich et al., 2018 |
| PBMC (hg19) | ATAC | Target | 7,828 | 452,004 | 10 | Ciero | Granja et al., 2019 |
| BMMC (hg19) | ATAC | Target | 14,753 | 452,004 | 13 | Ciero | Granja et al., 2019 |
| Breast cancer (hg19) | ATAC | Target | 11,116 | 155,403 | 7 | Signac | Kumegawa et al., 2022 |
| PBMC D10T1 (hg19) | ATAC | Reference/Target | 2,588 | 452,004 | 9 | Ciero | Granja et al., 2019 |
| PBMC D12T1,2,3 (hg19) | ATAC | Reference/Target | 3,070 | 452,004 | 9 | Ciero | Granja et al., 2019 |
